# Emergence of Sparse Coding, Balance and Decorrelation from a Biologically-Grounded Spiking Neural Network Model of Learning in the Primary Visual Cortex

**DOI:** 10.1101/2024.12.05.627100

**Authors:** Marko A. Ruslim, Martin J. Spencer, Hinze Hogendoorn, Hamish Meffin, Yanbo Lian, Anthony N. Burkitt

## Abstract

Many computational studies attempt to address the question of information representation in biological neural networks using an explicit optimization based on an objective function. These approaches begin with principles of information representation that are expected to be found in the network and from which learning rules can be derived.

This study approaches the question from the opposite direction; beginning with a model built upon the experimentally observed properties of neural responses, homeostasis, and synaptic plasticity. The known properties of information representation are then expected to emerge from this substrate.

A spiking neural model of the primary visual cortex (V1) was investigated. Populations of both inhibitory and excitatory leaky integrate-and-fire neurons with recurrent connections were provided with spiking input from simulated ON and OFF neurons of the lateral geniculate nucleus. This network was provided with natural image stimuli as input. All synapses underwent learning using spike-timing-dependent plasticity learning rules. A homeostatic rule adjusted the weights and thresholds of each neuron based on target homeostatic spiking rates and mean synaptic input values.

These experimentally grounded rules resulted in a number of the expected properties of information representation. The network showed a temporally sparse spike *response* to inputs and this was associated with a sparse *code* with Gabor-like receptive fields. The network was balanced at both slow and fast time scales; increased excitatory input was balanced by increased inhibition. This balance was associated with decorrelated firing that was observed as population sparseness. This population sparseness was both the cause and result of the decorrelation of receptive fields. These observed emergent properties (balance, temporal sparseness, population sparseness, and decorrelation) indicate that the network is implementing expected principles of information processing: efficient coding, information maximization (’infomax’), and a lateral or single-layer form of predictive coding.

These emergent features of the network were shown to be robust to randomized jitter of the values of key simulation parameters.

## 2 Introduction

Computational models of learning in the cortex are a powerful tool to integrate different domains of knowledge regarding brain function. Simulations can lead to a consistent understanding across qualitative and quantitative information, and can combine theoretical restrictions with parameters from experimental results. Many studies of biological neural networks begin ‘top-down’ with normative principles of information representation such as predictive coding (Rao and Ballard, 1999) or sparse coding (Olshausen and Field, 1997) and then derive the properties of an underlying neural network. This study takes a ‘bottom-up’ approach. A biologically grounded model was created based on experimental data of the properties of neural response and plasticity and analysis of the results used to determined whether useful properties of neural representation would emerge.

### 2.1 What are Biologically Grounded Neural Networks?

In this study we define a Biologically Grounded Neural Network (BGNN) to have the following experimentally observed features related to the network’s architecture, neural dynamics and learning dynamics:

- In the visual cortex input from separate populations of ON and OFF LGN neurons encoding local positive and negative differences in luminance relative to background (Ichinose and Habib, 2022) (Figure 1A).
- Distinct populations of excitatory and inhibitory neurons rather than individual neurons that are simultaneously excitatory and inhibitory; a requirement known as Dale’s Law (Eccles, 1976). (Figure 1B).
- Lateral connections between neurons within a layer of the network. These are not included in simulations based on objective functions (Figure 1B).
- Learning rules that use local information available at the synapse rather than any explicit global objective function used in most studies or a back-propagation approach used in artificial neural networks.
- Spikes to transmit and encode information rather than scalar rate values often used in neural networks.
- Spike-Timing-Dependent Plasticity (STDP) rules to adjust synaptic weights based on the timing differences of spikes that rate-based Hebbian rules ignore (Feldman, 2012). (Figure 1C).

**Figure 1:**
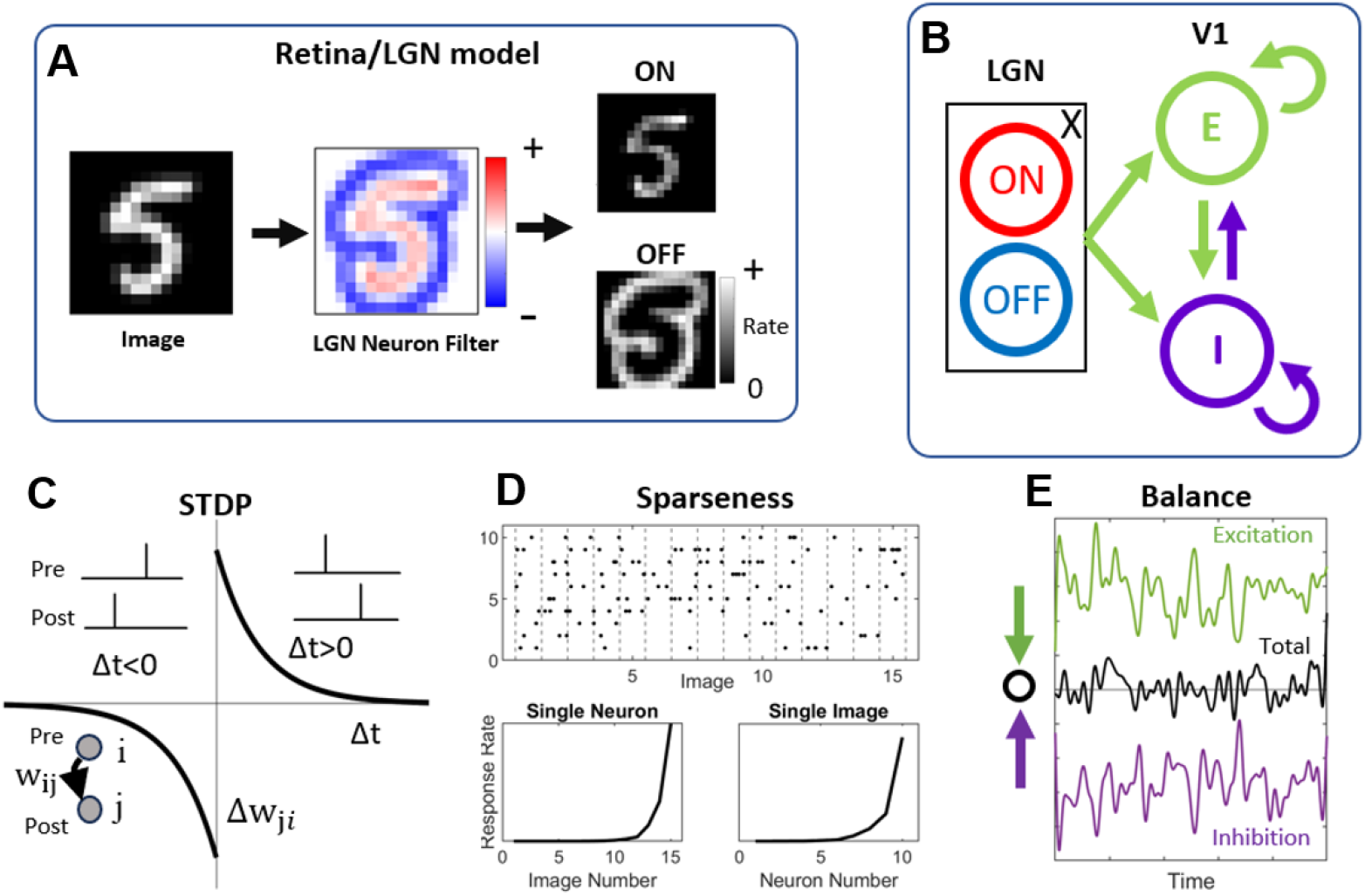
Features of Biological Neural Network Models: (A) Neurons of the Lateral Geniculate Nucleus are simulated as Poisson neurons responding at a rate that is proportional to a centre surround filter at each location of the visual field. (B) Separate populations of excitatory and inhibitory neurons received feed-forward input from the LGN neurons and have all-to-all lateral connectivity. (C) Synaptic weights between all populations are adjusted based on the relative timing of the input and output spikes. (D) Sparseness is quantified both as the response rate of each neuron across images (temporal sparseness), as well as the response of each neuron across the population in response to an individual image (population sparseness). (E) Balance is observed as a correlation in the magnitude of excitation and inhibition, which can occur at different time-scales.

Table 1 compares previous sparse coding models that learn V1 properties using some of the biologically-grounded features mentioned above. Previous models are missing in one or more of these features, putting into question how the brain can implement sparse coding. A model that is worth highlighting is by Brito and Gerstner (2024). Their model includes separate ON and OFF LGN inputs, spiking inputs and outputs, as well as STDP. However, their model does not include a separate inhibitory subpopulation and instead implements a single output population with recurrent inhibition. Additionally, their model does not capture the diverse RF shapes seen in biology that other sparse coding models can account for (Chauhan et al., 2021; King et al., 2013; Lian et al., 2019; Rehn and Sommer, 2007; Zylberberg et al., 2011). The sparse coding model by King et al. (2013) includes a separate inhibitory subpopulation, but omit connections such as the feedforward connection to inhibitory neurons and recurrent excitatory connections that are seen in V1. The model presented in this work incorporates these features of LGN ON/OFF separation, spiking neurons, learning via STDP and a separate inhibitory subpopulation.

**Table 1:**
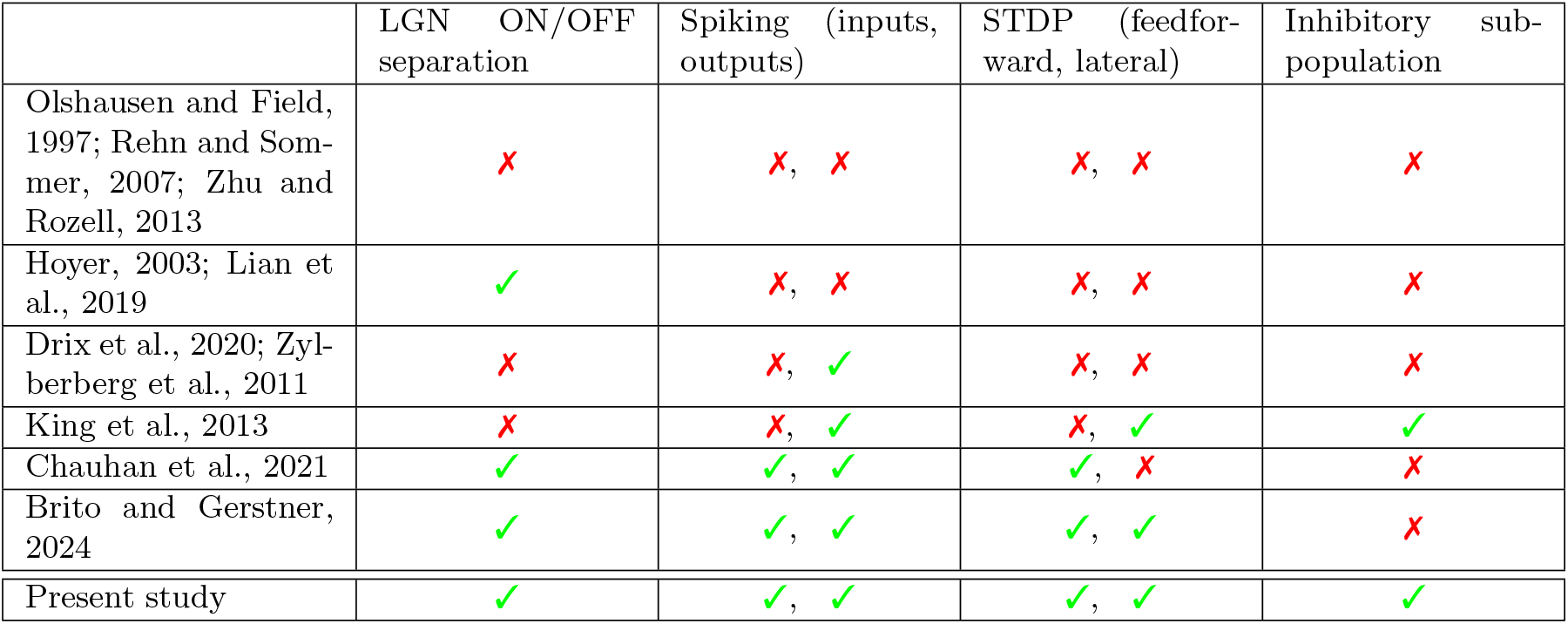
Comparison of sparse coding models that learn V1 properties. Row headings are sparse coding models or models based on sparse coding and column headings are features of biologically-grounded models.

|  | LGN ON/OFF separation | Spiking (inputs, outputs) | STDP (feedforward, lateral) | Inhibitory sub-population |
| --- | --- | --- | --- | --- |
| Olshausen and Field, 1997; Rehn and Sommer, 2007; Zhu and Rozell, 2013 | ✗ | ✗, ✗ | ✗, ✗ | ✗ |
| Hoyer, 2003; Lian et al., 2019 | ✓ | ✗, ✗ | ✗, ✗ | ✗ |
| Drix et al., 2020; Zylberberg et al., 2011 | ✗ | ✗, ✓ | ✗, ✗ | ✗ |
| King et al., 2013 | ✗ | ✗, ✓ | ✗, ✓ | ✓ |
| Chauhan et al., 2021 | ✓ | ✓, ✓ | ✓, ✗ | ✗ |
| Brito and Gerstner, 2024 | ✓ | ✓, ✓ | ✓, ✓ | ✗ |
| Present study | ✓ | ✓, ✓ | ✓, ✓ | ✓ |

### 2.2 Sparseness, Balance, and Decorrelation

In addition to these underlying biological properties of the network and its dynamics, there are emergent properties that relate to network stability, energy efficiency, and optimum information representation. Table 2 compares the emergent properties of these sparse coding models with the present study.

**Table 2:** Comparison of the emergent properties of V1 sparse coding models. Row headings are sparse coding models or models based on sparse coding and column headings are emergent features. Asterisks on the cited papers indicate those that also include an *n*_*x*_ vs *n*_*y*_ plot to illustrate the diversity of receptive field shapes.

|  | Sparseness |  | Balance | E-I Decorrelation |
| --- | --- | --- | --- | --- |
|  | Imposed | Emerged | Analysed |  |
| Olshausen and Field, 1997; Hoyer, 2003; Rehn and Sommer, 2007*; Zhu and Rozell, 2013; Lian et al., 2019* | ✓ |  | ✗ | ✗ |
| Zylberberg et al. 2011*; Drix et al. 2020; Chauhan et al. 2021*; Brito and Gerstner 2024 |  | ✓ | ✗ | ✗ |
| King et al. 2014 |  | ✓ | ✗ | ✓ |
| Present study* |  | ✓ | ✓ | ✓ |

Neurons in the cortex are observed to show a **sparse response** to sensory stimuli; individual neurons respond strongly to only a minority of stimuli (temporal sparseness) and an individual stimulus results in activity in only a minority of neurons (population sparseness) (Figure 1D). This sparse response of the cortex is associated with a **sparse code** that achieves an energy efficient representation of the input data, and maps the incoming sensory data to a small set of causes. These causes are the receptive fields of each neuron; the particular visual input that causes a neuron to respond. In the visual cortex most spatial receptive fields are experimentally observed to be well fitted by a Gabor function (Ringach, 2002), a result that matches the theoretical predictions of sparse-coding models (Bell and Sejnowski, 1997; Olshausen and Field, 1996; Rehn and Sommer, 2007; Zhu and Rozell, 2013)).

Biological networks also exhibit **balance** between excitation and inhibition (Barral and D Reyes, 2016; Froemke, 2015; Haider et al., 2006). Balance is a dynamical feature of biological neural networks and is known to enforce stability and prevent runaway pathological responses (van Vreeswijk and Sompolinsky, 1996). Balance can enhance the precision of cortical representations (Boerlin et al., 2013). The degree of balance can be evaluated at a range of timescales, from ‘loose’ balance for long time scales, to ‘tight’ balance for fine time scales (Denève and Machens, 2016).

Loose balanced is a result of the overall magnitude of excitatory and inhibitory input to the neurons of the network, where total input is due to a combination of the relative number of excitatory and inhibitory neurons in the network, their relative firing rates, and their total synaptic weight.

Tight balance can be observed in temporal correlations on the scale of milliseconds between the rapid changes in the excitation and inhibition provided to individual neurons, as illustrated in Figure 1E.

Balance also appears to be associated with the decorrelation of responses between neurons; if inhibition grows to match excitation then the neuron will be prevented from firing. This in turn leads to decorrelation of the receptive fields of each neuron. Effectively this creates competition between the neurons in the network.

Neurons may have correlated spike rates over time and/or correlated receptive fields. **Decorrelation** in biological neural networks refers to a tendency for neurons to compete and reduce redundancy in the network’s representation of sensory input. The population sparseness described above is an example of de-correlated firing between individual neurons (Tetzlaff et al., 2012).

Via plasticity of the weights between the LGN neurons and V1 neurons competition to produce spikes is the cause of diversity in the set of receptive fields, however de-correlated firing is also the result of this diversity in receptive fields. This diversity minimizes redundancy in the network’s representation of features of the visual field.

### 2.3 Proposed Model

The model in this investigation used a BGNN and allowed the the properties of sparseness, balance, and decorrelation to emerge. By creating a BGNN model of the visual cortex, including an accurate model of STDP, it was possible to make a convincing bottom-up connection between biological processes and these principles of information representation in the brain.

### 2.4 Application in Artificial Neural Networks

An additional purpose in simulating learning in BGNNs is as a biomimetic approach to improving Artificial Neural Networks (ANNs). In this context the model proposed in this study of a BGNN can be thought of as a specific form of an Artificial Spiking Neural Network (ASNN) (Rathi et al., 2023).

Existing approaches to ANNs use an extremely large amount of energy during training (Desislavov et al., 2023; Gelles, 2024). When implemented in neuromorphic hardware ASNNs are orders of magnitude more energy efficient (Basu et al., 2018). In addition, ASNNs may have advantages over conventional ANNs in temporal event-based information representation (Afshar et al., 2020). ASNNs may be trained using a ‘shadow’ ANN before implementation in a SNN (Rueckauer et al., 2017), or back-propagation learning is adapted for use in a SNN (Eshraghian et al., 2023; Neftci et al., 2019). While there has been moderate success using these methods, neither have matched the success of conventional ANNs and neither take full advantage of low-energy usage neuromorphic hardware.

A secondary goal of this study is to create a BGNN that could in the future provide comparable computational performance to ANNs while also being possible to implement in neuromorphic hardware that has lower energy usage than existing computational systems.

## 3 Methods

### 3.1 Network Architecture

The network simulated in this study consisted of *N* ^E^ = 400 excitatory (E) and *N* ^I^ = 100 inhibitory (I) neurons representing model V1 simple cells. These were modelled as Leaky Integrate-and-Fire (LIF) neurons (Burkitt, 2006). LIF models can accurately capture many of the most salient properties of neuron in the brain such as the temporal integration of synaptic inputs and a non-linear (or threshold) firing mechanism (Mensi et al., 2016). Most importantly, the LIF model produces action potentials rather than describing the neural activity as a scalar firing rate. Among these model V1 neurons there was all-to-all lateral connectivity with no self-connections. This is a reasonable assumption in simulating a small region of the cortex of the scale, for example, of a cortical column.

The model V1 neurons received feedforward thalamic input from a population (X) of *N* ^X^ = 512 Lateral Geniculate Nucleus (LGN) neurons. These LGN neurons were modelled as linear-nonlinear Poisson neurons that process the visual stimuli. The LGN input was split into 256 ON and 256 OFF sub-populations that each respond to light increments or light decrements (respectively) in their central receptive field, relative to background levels (Ichinose and Habib, 2022).

The visual stimuli to the thalamic neurons was provided by images with 16*×*16 = 256 pixels. All synapses are plastic and are trained during learning.

### 3.2 LGN Processing

Raw images from the van Hateren dataset (van Hateren and van der Schaaf, 1998) were used to train the model under STDP. This dataset contains over 4000 images, each containing a natural scene. For these simulations, 1000 images were randomly chosen. Before presenting these images to the network, these images were first filtered using a centre-surround receptive field filter in the linear-nonlinear model neuron. This mimics the processing performed by the retina and LGN. The filter is a divisively normalised difference-of-Gaussians (Ratliff et al., 2010):

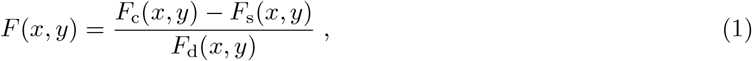

where *F* (*x, y*) is the intensity at image location (*x, y*) after applying the spatial filtering, and *F*_c_, *F*_s_ and *F*_d_ are normalized, concentric, isotropic Gaussian filters with standard deviations *σ*_c_ = 1, *σ*_s_ = 1.5, and *σ*_d_ = 1.5, for center, surround and divisive normalization respectively.

Random patches of size 16 *×* 16 were sampled from these LGN-processed images, and selected as the input stimuli to the network. The positive and negative values were separated into ON and OFF input sub-populations. Just as is the case in the neurons of the retina and LGN, the values are scaled to give appropriate firing rates across a large number of images. In particular image patches were scaled by a factor of *c*_r_ = 70 to yield suitable firing rates (i.e., mean input firing rates of 20 Hz). Spikes were generated by simulating Poisson processes. To simulate the saturation of retinal and LGN neurons, the Poisson firing rates of the LGN neurons were bounded at 100 Hz though actual generated spike rates generated by the Poisson process can exceed this.

### 3.3 V1 Neuron Dynamics

The neuron model used in the simulations is a current-based LIF neuron (Burkitt, 2006). The governing dynamics for the membrane potential, 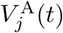, of neuron *j* in the output population, A, where A ∈ {E, I} represents the excitatory and inhibitory populations, is:

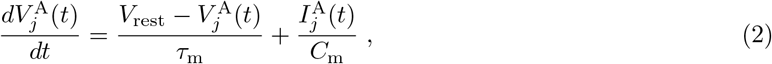

where *C*_m_ is the membrane capacitance, *τ*_m_ is the passive membrane time constant, and *V*_rest_ is the resting membrane potential, which is assumed here to be the same for both populations of neurons. When the membrane potential crosses an adaptive spiking threshold, 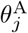, an action potential is generated and the membrane potential is reset to the resting potential. Additionally there is a lower bound for the membrane potential at *V*_min_, analogous to the inhibitory reversal potential. The synaptic input, 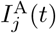, to neuron *j* in neural population A is modelled as an instantaneous current injection (for notational convenience the membrane capacitance parameter, *C*_m_, is henceforth absorbed into the scaling of the weight matrix, **W**^AB^, of the network):

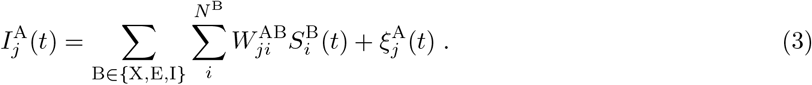

The synaptic input consists of the feed-forward input from the LGN population X, recurrent inputs from other cortical neurons and Gaussian white noise, 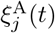, mimicking additional inputs from other more distant cortical neurons. The synaptic weight from presynaptic neuron *i* = 1, 2, …, *N* ^B^ in population B to postsynaptic neuron *j* in population A is 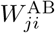. The spike train, 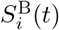, of neuron *i* in population B is represented by the sum of Dirac delta functions at the spike times. Gaussian white noise is added to the current at every *δt* = 1 ms timestep sampled from the distribution 𝒩 (*µ*_*ξ*_, *σ*_*ξ*_^2^), where *µ*_*ξ*_ = 0 and *σ*_*ξ*_ = *δt* = 1. The value of *σ*_*ξ*_ was chosen to have a small effect, namely increasing stochasticity of output spikes, but not a dominant effect, since synaptic strength and connection probability decays with distance (Perin et al., 2011).

### 3.4 STDP and Homeostasis

#### 3.4.1 STDP

The learning rules in this network utilize Spike-Timing-Dependent Plasticity (STDP). Experimentally, the form of STDP depends on the type of synapse. For the synapses between two excitatory neurons, that is, each of the entries in the weight matrices **W**^EX^ and **W**^EE^, the plasticity rule used in this study is the minimal triplet STDP rule, which has been shown to fit experimental data for pyramidal neurons of the rat visual cortex and hippocampus well (Pfister and Gerstner, 2006). This STDP window function is asymmetric and has an adaptation to the spike output rate. Learning is triggered whenever the postsynaptic neuron *j* spikes (at time *t*_*j*_) or the presynaptic neuron *i* spikes (at time *t*_*i*_):

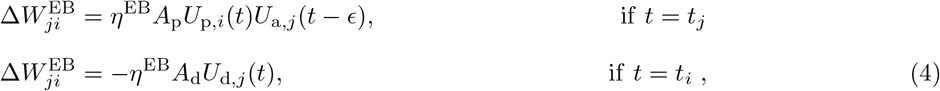

where 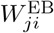 represents the synaptic weight from neuron *i* in population B ∈ {X, E} to neuron *j, η*^EB^ is the learning rate and *ϵ* is a small positive constant to ensure that the weight change is updated before the synaptic trace variable, *U*_*α,i*_(*t*), is updated. This synaptic trace is the convolution of a truncated exponential kernel, *K*_*α*_(*t*), with the neuron’s spike train:

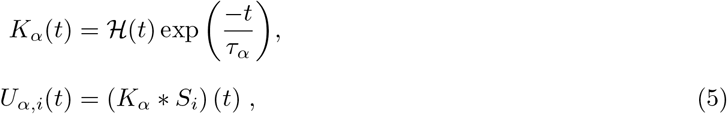

where *H*(*t*) is the Heaviside step function and *τ*_*α*_ are the STDP time constants for *α* = p, d, a. The STDP amplitudes are *A*_p_ and *A*_d_, which are the potentiation and depression sides of the triplet STDP respectively. The STDP time constants are *τ*_p_ and *τ*_d_. The third STDP time constant, *τ*_a_, is a long time constant that essentially calculates a moving average of the postsynaptic spike train. This feature distinguishes the triplet model from classical STDP, which only considers presynaptic and postsynaptic spike pairs.

It has been shown that the minimal triplet model can be mapped to the BCM learning rule by averaging over spike times to give a rate-based description if presynaptic and postsynaptic spikes are uncorrelated (Bienenstock et al., 1982; Pfister and Gerstner, 2006). The equivalent threshold in BCM rule, *ϕ*, between potentiation and depression in this minimal triplet model is given by the firing rate *ϕ* defined as:

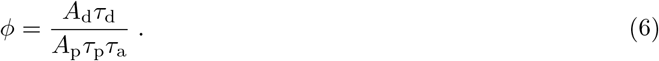

The parameter *ϕ* is chosen to be the same as the target firing rate of the excitatory neurons, *ϕ* = *ρ*^E^. This target firing rate is also maintained through homeostatic plasticity (Section 3.4.3). The triplet STDP parameters, namely the potentiation and depression coefficient values, *A*_p_ and *A*_d_, and triplet STDP time constants are chosen to achieve this value for *ϕ*. Setting the STDP parameters in this way achieves rate equilibrium for uncorrelated input and output spikes. In the original BCM rule, the threshold varies to stabilize learning, and although the threshold in this triplet rule is fixed, the adaptive spiking threshold and weight normalization may take this role.

For all other synapses, namely each of the entries in the weight matrices **W**^EI^, **W**^IX^, **W**^IE^, and **W**^II^, the plasticity rule is a purely symmetrical STDP rule, which is similarly activated when the presynaptic or postsynaptic neuron fires a spike. This corresponds closely to rate-based Hebbian learning and has been observed experimentally for inhibitory synapses of mouse auditory cortex (D’amour and Froemke, 2015). This rule was also adopted for excitatory to inhibitory synapses as a simplifying assumption. The symmetrical STDP rule is in the form:

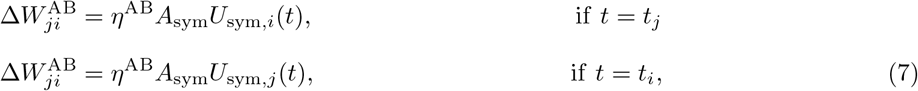

which has an STDP time constant of *τ*_sym_. Note that although inhibitory synapses have negative weights, the symmetrical STDP rule acts to strengthen these weights.

#### 3.4.2 Weight Normalization

All synaptic weights to a neuron have an L1 norm upper bound. If the L1 norm is exceeded due to STDP during training, then those weights are normalised back to their upper bound through subtractive normalisation. For example, 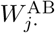, which is the weights from population B ∈ {E, I, X} to postsynaptic neuron *j* in population A ∈ {E, I}, undergoes subtractive normalisation if the L1 norm exceeds 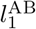:

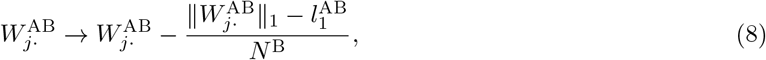

where 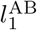 is the upper bound of the L1 norm. There is also a constraint of non-negative weights: 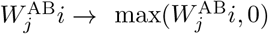 which is enforced to obey Dale’s law. However, due to this constraint, the L1 norm of the weights may still exceed the upper bound. Therefore, multiplicative normalisation is also applied:

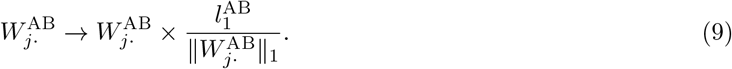

Weight normalisation is a necessary feature to ensure stability of weights and consequently, spiking activity in the network. Weight normalisation is akin to activity-dependent synaptic scaling, which is a homeostatic process that acts to keep firing activity within stable levels (Turrigiano et al., 1998).

#### 3.4.3 Homeostatic Plasticity

Homeostatic plasticity is also employed in the form of an adaptive spiking threshold, 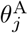, for neuron *j* in population A = {E, I} (Földiák, 1990; Zylberberg et al., 2011):

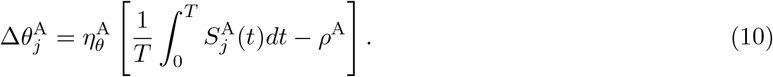

This homeostatic rule ensures the excitatory (inhibitory) neurons have a time-average firing rate of *ρ*^E^ (*ρ*^I^), by adjusting the spiking threshold of the LIF dynamics. An upper bound is enforced to the change in spiking threshold, being 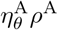. An adaptive spiking threshold is a mechanism of intrinsic plasticity that affects the excitability of a neuron, though other mechanisms of intrinsic neuronal excitability exist (Debanne et al., 2019).

### 3.5 Synaptic Balance in the Network

Balanced network theory was used to determine appropriate mean weights and inform the choice of L1 upper bounds of synaptic weights. The idea behind the theory of balanced networks is that excitatory and inhibitory inputs to a neuron are balanced, and that spiking activity is irregular and driven by fluctuations. The balance condition under large *N*, which is the total number of neurons in the network, is that the mean total synaptic current to a neuron scales as *O*(1). Balance and a stable solution is obtained if:

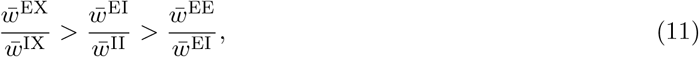

where 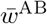 is the mean synaptic weight from neurons in population B to neurons in population A. This relation is similar to the balanced condition derived in other studies (Gu et al., 2018; Tian et al., 2020; van Vreeswijk and Sompolinsky, 1998). Although balance theory was used as a guiding principle to set the upper bounds of the L1 weight norm, it is important to note that this balance condition was derived under specific conditions to apply the mean-field theory: namely that input neurons fire at a constant and identical firing rate, and that output neurons are independent due to the random structure of the weights. However, in our simulations, input neurons fire at different firing rates depending on the visual stimuli, and the weights in our network show structure due to learning.

Another consideration is that the ratio of the number of excitatory neurons to inhibitory neurons is approximately 4:1 in biology (DeFelipe and Fariñas, 1992). As such, there should be compensatory features to ensure approximate balance between excitatory and inhibitory inputs. The compensatory features used in our simulations are that the target firing rate of inhibitory neurons is double the target firing rate of excitatory neurons, and that the average synaptic strength of inhibitory synapses is roughly double that of lateral excitatory connections.

The network consists of an excitatory to inhibitory neuron ratio of 4:1. To establish and maintain loose balance, the target firing rate of the inhibitory neurons is double the firing rate of the excitatory neurons and the average synaptic strength of inhibitory synapses is roughly double that of excitatory synapses:

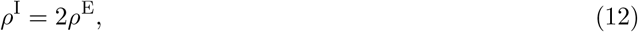

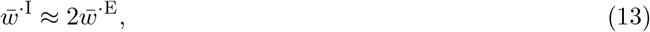

where 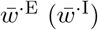 is the mean synaptic weight with an excitatory (inhibitory) presynaptic neuron. The values of the L1 weight norm upper bounds were chosen to adhere to Equations 11 and 13 and are shown in Table 3. Equations 12 and 13 are enforced by the adaptive spiking threshold (Equation 10) and homeostatic weight normalization (Equations 8 and 9).

**Table 3:**
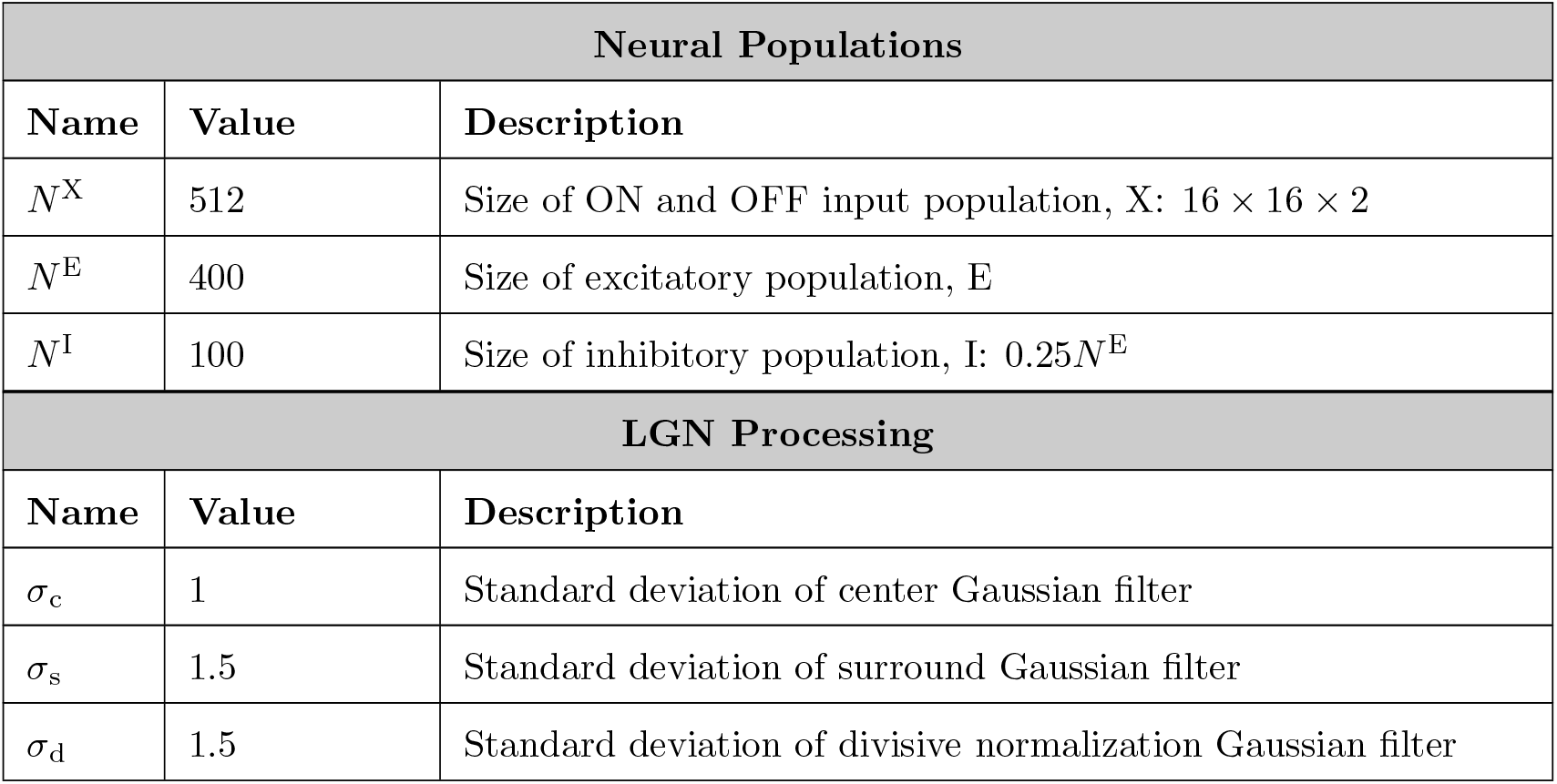
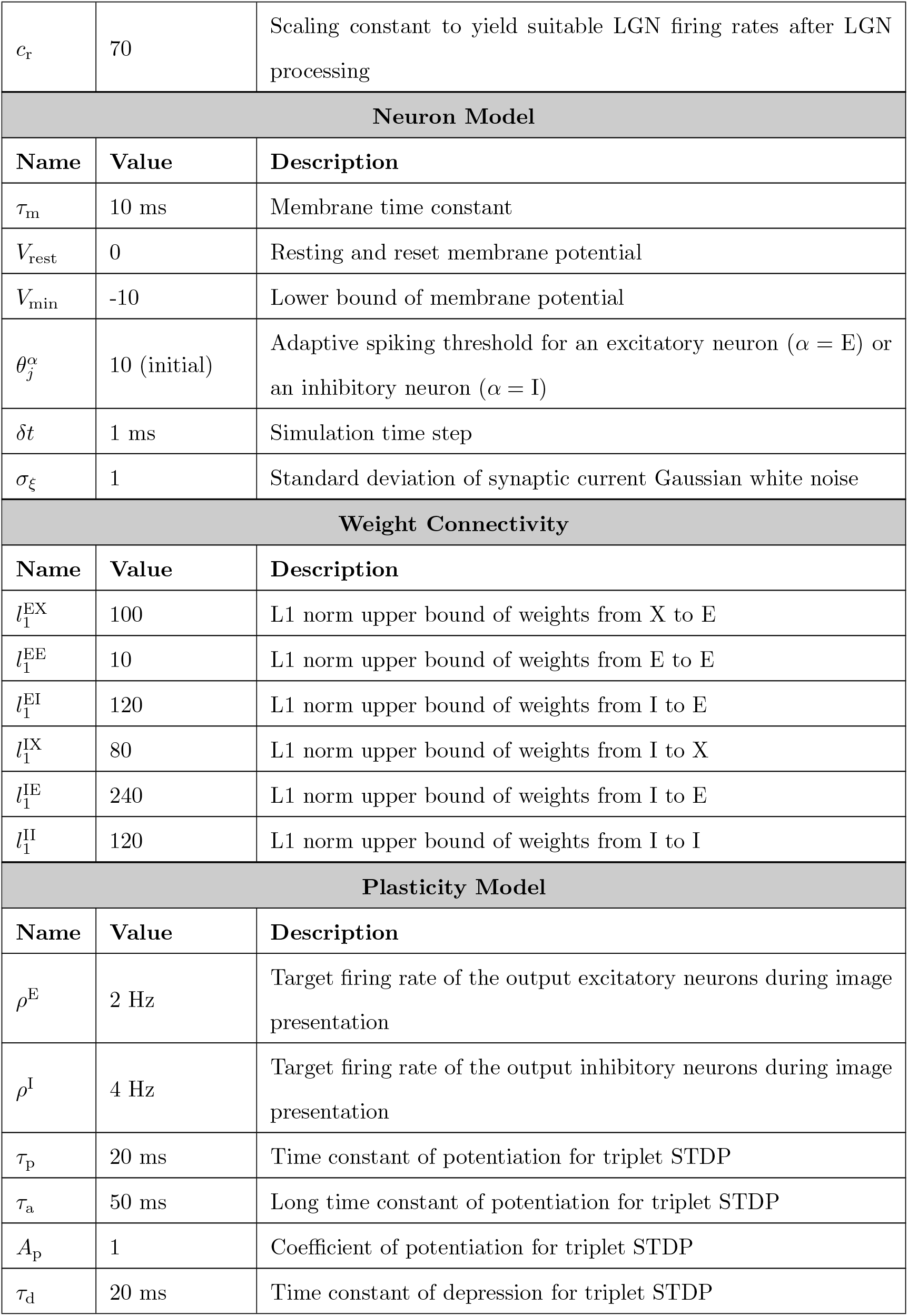
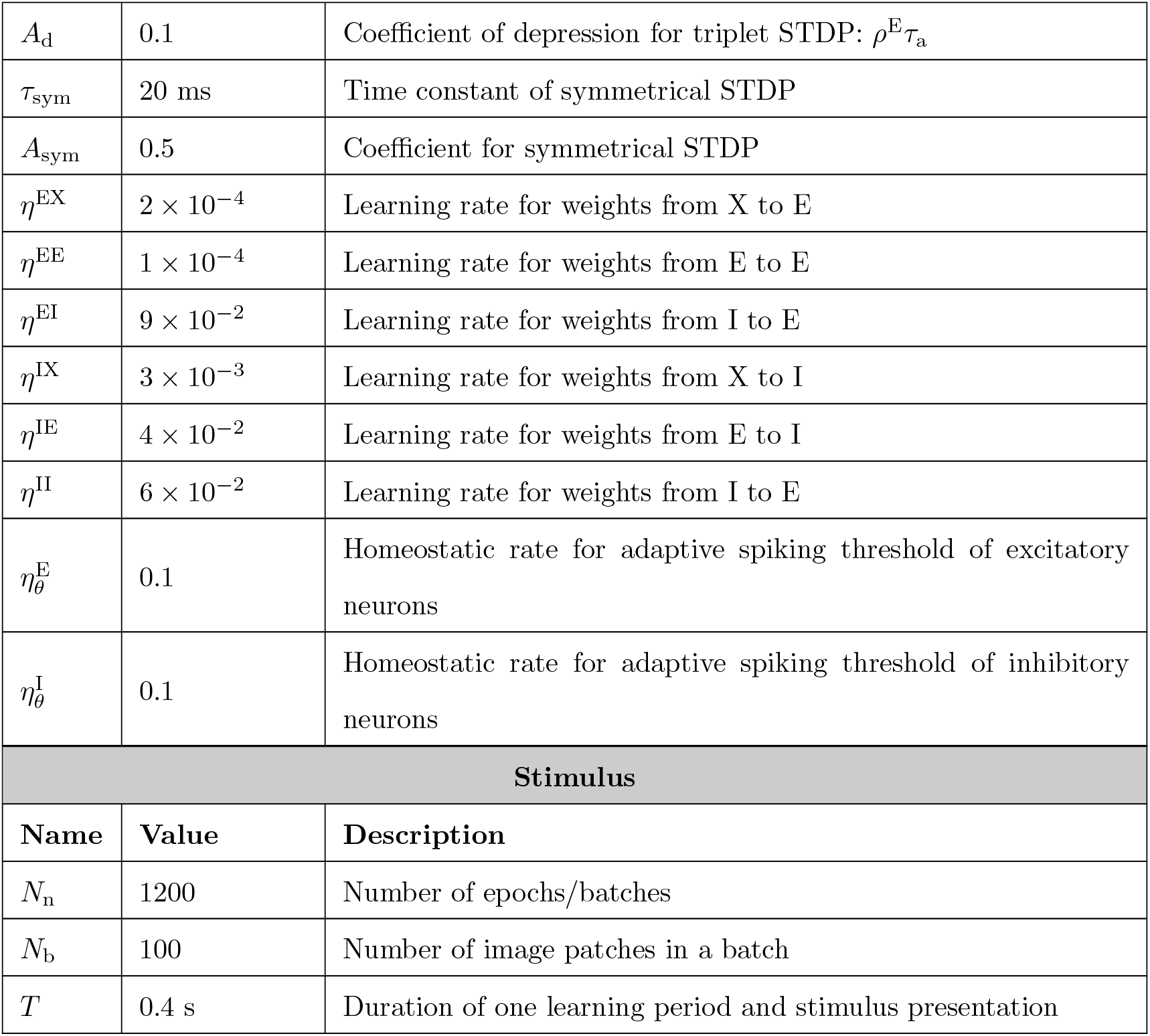
Simulation parameter summary for the network model.

| Neural Populations |  |  |
| --- | --- | --- |
| Name | Value | Description |
| $N^X$ | 512 | Size of ON and OFF input population, X: $16 \times 16 \times 2$ |
| $N^E$ | 400 | Size of excitatory population, E |
| $N^I$ | 100 | Size of inhibitory population, I: $0.25N^E$ |
| LGN Processing |  |  |
| Name | Value | Description |
| $\sigma_c$ | 1 | Standard deviation of center Gaussian filter |
| $\sigma_s$ | 1.5 | Standard deviation of surround Gaussian filter |
| $\sigma_d$ | 1.5 | Standard deviation of divisive normalization Gaussian filter |
| $c_r$ | 70 | Scaling constant to yield suitable LGN firing rates after LGN processing |
| **Neuron Model** |  |  |
| **Name** | **Value** | **Description** |
| $\tau_m$ | 10 ms | Membrane time constant |
| $V_{\text{rest}}$ | 0 | Resting and reset membrane potential |
| $V_{\text{min}}$ | -10 | Lower bound of membrane potential |
| $\theta_j^\alpha$ | 10 (initial) | Adaptive spiking threshold for an excitatory neuron ($\alpha = \text{E}$) or an inhibitory neuron ($\alpha = \text{I}$) |
| $\delta t$ | 1 ms | Simulation time step |
| $\sigma_\xi$ | 1 | Standard deviation of synaptic current Gaussian white noise |
| **Weight Connectivity** |  |  |
| **Name** | **Value** | **Description** |
| $l_1^{\text{EX}}$ | 100 | L1 norm upper bound of weights from X to E |
| $l_1^{\text{EE}}$ | 10 | L1 norm upper bound of weights from E to E |
| $l_1^{\text{EI}}$ | 120 | L1 norm upper bound of weights from I to E |
| $l_1^{\text{IX}}$ | 80 | L1 norm upper bound of weights from I to X |
| $l_1^{\text{IE}}$ | 240 | L1 norm upper bound of weights from I to E |
| $l_1^{\text{II}}$ | 120 | L1 norm upper bound of weights from I to I |
| **Plasticity Model** |  |  |
| **Name** | **Value** | **Description** |
| $\rho^{\text{E}}$ | 2 Hz | Target firing rate of the output excitatory neurons during image presentation |
| $\rho^{\text{I}}$ | 4 Hz | Target firing rate of the output inhibitory neurons during image presentation |
| $\tau_p$ | 20 ms | Time constant of potentiation for triplet STDP |
| $\tau_a$ | 50 ms | Long time constant of potentiation for triplet STDP |
| $A_p$ | 1 | Coefficient of potentiation for triplet STDP |
| $\tau_d$ | 20 ms | Time constant of depression for triplet STDP |
| $A_d$ | 0.1 | Coefficient of depression for triplet STDP: $\rho^E \tau_a$ |
| $\tau_{\text{sym}}$ | 20 ms | Time constant of symmetrical STDP |
| $A_{\text{sym}}$ | 0.5 | Coefficient for symmetrical STDP |
| $\eta^{\text{EX}}$ | $2 \times 10^{-4}$ | Learning rate for weights from X to E |
| $\eta^{\text{EE}}$ | $1 \times 10^{-4}$ | Learning rate for weights from E to E |
| $\eta^{\text{EI}}$ | $9 \times 10^{-2}$ | Learning rate for weights from I to E |
| $\eta^{\text{IX}}$ | $3 \times 10^{-3}$ | Learning rate for weights from X to I |
| $\eta^{\text{IE}}$ | $4 \times 10^{-2}$ | Learning rate for weights from E to I |
| $\eta^{\text{II}}$ | $6 \times 10^{-2}$ | Learning rate for weights from I to E |
| $\eta_{\theta}^{\text{E}}$ | 0.1 | Homeostatic rate for adaptive spiking threshold of excitatory neurons |
| $\eta_{\theta}^{\text{I}}$ | 0.1 | Homeostatic rate for adaptive spiking threshold of inhibitory neurons |
| **Stimulus** |  |  |
| **Name** | **Value** | **Description** |
| $N_n$ | 1200 | Number of epochs/batches |
| $N_b$ | 100 | Number of image patches in a batch |
| $T$ | 0.4 s | Duration of one learning period and stimulus presentation |

### 3.6 Training

The initial weight connectivity structure was random and sparse. Some weights were initialised to high values whereas others were initialised to low values:

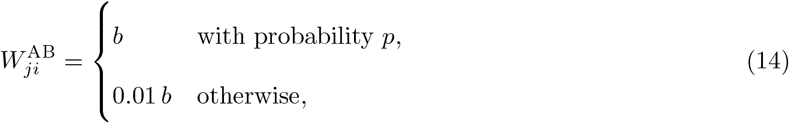

where *b* is sampled from a normal distribution with mean 1 and standard deviation 0.5, and *p* = 0.2 is the probability of a strong synaptic connection. This method of initialization is similar to how balanced networks are initialized (Brunel, 2000; Gu et al., 2018; Tian et al., 2020; van Vreeswijk and Sompolinsky, 1998). In our simulations, some additional jitter is introduced. However, other methods of initialisation such as sampling weight values from an exponential distribution lead to similar results. Following this, the weights are multiplicatively scaled to have their L1 norm equal to its upper bound (Equation 9). It was observed that when weights were initialised with very small non-zero values, the L1 weight norms all approached upper bound, and took much longer to learn (Supplementary Material 6).

The network is trained on static natural images. Each batch, which consists of *N*_b_ = 100 image patches was presented for *T* = 400 ms in parallel. The output spikes of model V1 neurons were generated as described in Section 3.3 with sub-threshold LIF membrane dynamics given by Equation 2. This presentation time gives the network enough time to respond to the stimulus and is longer than the STDP time constants. Before training, the spiking thresholds were allowed to reach stable values by running the network with an adaptive spiking threshold but no weight plasticity for 100 batches (100 *×* 100 *×* 400 ms ≈ 67 mins in model time). During training, the weights are updated through the weight learning rules (Equations 4, 7, 8, 9), as well as the spiking threshold through the homeostatic rule (Equations 10). Training occurs over *N*_*n*_ = 1200 batches (1200 *×* 100 *×* 400 ms ≈ 11.1 hrs model time). The weights appeared to show structure and converge with this number of batches. At batch number 0.3*N*_n_ and 0.6*N*_n_, the learning and homeostatic rates, *η*^AB^ and 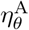 were divided by a factor of 2, in order to reduce fluctuations in weight dynamics and ensure stable convergence. However, the results are not sensitive to slight changes in batch number or learning rates. We also investigated if baseline firing to background input before and after learning is maintained, even with the learnt weight structure. This is shown in Supplementary Information 6

### 3.7 Code Availability

The code of implementing the model and running the simulation was written in MATLAB (R2023a), and is available at: https://github.com/marko-ruslim/bio-spiking-v1-learning.

### 3.8 Tabular Summary of Parameters

### 3.9 Analysis Methods

#### 3.9.1 Receptive Field Analysis

The receptive field refers to the ideal sensory stimulus that will trigger the firing of a neuron. Classically, the receptive field is measured in experiments by presenting visual stimuli such as spots of light, bars or gratings, and then measuring the neuron’s response. White noise stimulation has also been used as the visual stimuli and this provides an unbiased estimate of the receptive field (Chichilnisky, 2001; Schwartz et al., 2006). Properties of the visual neuron can be inferred from their receptive field such as orientation and spatial frequency tuning. Similar to experiments, a spike-triggered average was performed and Gabor-function fitting to the model simple cells.

##### Spike-Triggered Average

Following training, the receptive fields (RFs) of the modelled V1 cells are estimated by Spike-Triggered Average (STA) (Chichilnisky, 2001), also referred to as reverse correlation or white-noise analysis. Spatial Gaussian white noise stimulus with unit variance, **n**, is put through the same LGN pipeline, scaled by a factor of 2*c*_r_ to convert to spiking rate and presented to the network, and the firing rate, *r*, of a model cell is recorded. In order to convert Gaussian white noise to suitable pixel values between 0 and 255, the following expression is used: 50**n** + 100, with lower and upper bounds of 0 and 255 respectively. Its receptive field, **F**, can be estimated through a weighted average:

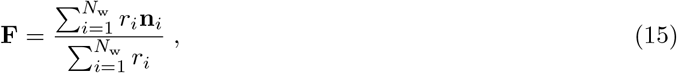

where *N*_w_ = 10^5^ is the number of white noise stimuli. The STA provides an unbiased estimate of a neuron’s receptive field only if the stimulus distribution is spherically symmetric, e.g., Gaussian white noise (Chichilnisky, 2001; Schwartz et al., 2006).

The results of the STA are up-sampled from 16 *×* 16 pixels to 160 *×* 160 pixels using bilinear interpolation.

##### Fitting Gabor-functions to receptive fields

After the RFs of the V1 neurons in the model have been characterized by STA, they were then fit with Gabor filters, similar to that done in previous studies using both experimentally recorded RFs (Ringach, 2002) and simulated RFs (Lian et al., 2019), which facilitates comparison of simulation results with experimental data. A 2D Gabor function, *G*(*x, y*), is defined as a sinusoidal plane wave multipled by a 2D Gaussian window. Similar to Lian et al. (2019), the fitting error is defined as the ratio of the sum of the squared residuals over that of the receptive field. The receptive fields that had a fitting error of less than 10% were described as well-fit to a Gabor function. A scatter plot of *n*_*x*_ = *σ*_*x*_*f*_*s*_ vs *n*_*y*_ = *σ*_*y*_*f*_*s*_ is constructed, which gives the distribution of the Gabor filter properties for both the model and experimental V1 neurons (Ringach, 2002.

#### 3.9.2 Quantifying Sparseness

As described in the introduction, neurons in the visual cortex are known to exhibit a sparse spike response to visual stimuli.

The modified Treves-Rolls sparseness metric was used to quantify **temporal** sparseness (sometimes called lifetime sparseness) for a given neuron (Willmore and Tolhurst, 2001):

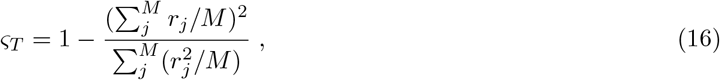

where *r*_*j*_ is the neuron’s response rate to image *j*, and *M* is the number of images used.

An identical metric was used to quantify the **population** sparseness in response to a given image:

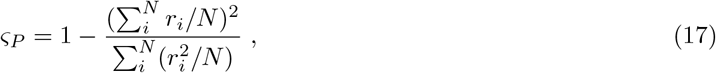

where *r*_*i*_ is each neuron *i*’s response rate to the image, and *N* is the number of neurons in the population.

The value of *ς* in each case lies between [0, 1] and approaches 1 for a highly sparse responses.

The temporal sparseness in Equation 16 is only for a single neuron and the population sparseness in equation 17 is only for a single image. To obtain a measure of the overall temporal and population sparseness the average value across neurons is used for temporal sparseness 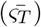 and the average value across images is used for population sparseness 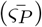.

#### 3.9.3 Quantifying Balanced Activity

As described in the introduction, experimental data from the cortex reveals that excitatory and inhibitory inputs to neurons are balanced over both slow and fast time-scales.

To examine balance over slow time scales (loose balance) the excitatory and inhibitory inputs to a neuron was examined on an image-by-image basis. An excitatory neuron was randomly selected. Its excitatory and inhibitory input was compared under conditions of both low and high feedforward excitatory inputs as described in the results. This was repeated for 40 neurons plotting the mean excitatory and total input as a function of the level of inhibitory input across images.

To examine balance over the fast time scales (tight balance) the cross-correlation between the excitatory and inhibitory inputs was used. This does not quantify the absolute ratio of excitation to inhibition but allows quantification of the time scale over which fluctuations in excitation are countered by fluctuations in inhibition.

#### 3.9.4 Quantifying Decorrelation

To discover the presence of a decorrelation effect in the network comparisons were made between the receptive fields of pairs of neurons. This was done using the correlation coefficient between their two STAs. The synaptic weight from one neuron to the other was then plotted as a function of this correlation coefficient. If the network has learned because pairs of neurons with correlated receptive fields might be expected to have an inhibitory effect on each other.

#### 3.9.5 Network Robustness

To test robustness of the results to parameter changes, Monte Carlo simulations were run in which the upper bounds for the L1 norm of the weights for E, I and X inputs were each varied. For each simulation, each of these parameters was scaled by a random number sampled from the uniform distribution between 1 − *c* and 1 + *c* for each postsynaptic neuron. For instance, 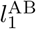 would become a *N* ^A^ *×* 1 vector where neurons in population A may have different weight norm upper bounds. Ten simulations were run where *c* = *{*0%, 10%, 20%, 40%*}*. To evaluate the performance of the network, three measures were used: the root-mean-squared (RMS) of the Pearson correlation coefficient between the receptive fields of all excitatory neuron pairs, and the temporal and population sparseness. The first measure quantifies the degree of correlation between excitatory receptive fields. It is calculated by first computing the Pearson correlation coefficient between the receptive fields of all excitatory cell pairs, then computing the RMS (King et al., 2013).

Simulation parameters remain the same (Table 3) except for the L1 norms which are jittered and the learning rates. Due to different L1 norms, the learning rates for each group of weights were adjusted to compensate. For example, for the weights from feedforward to excitatory neurons:

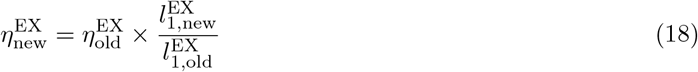

#### 3.9.6 Target Firing Rate and Network Size

Simulations with different target firing rates were run. In particular, excitatory neurons had target firing rate of 1 Hz, 2 Hz, 5 Hz, 10 Hz or 20 Hz. The inhibitory neurons had a target firing rate of double that of the excitatory neurons, namely 2 Hz, 4 Hz, 10 Hz, 20 Hz and 40 Hz, respectively. Ten simulations for each of these target firing rates were run.

Simulation parameters remain the same (Table 3) except for the target firing rates, the depression coefficient of triplet STDP and the learning rates. In order to satisfy equation 6, the coefficient of depression becomes: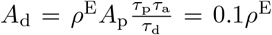. Additionally, due to different firing rates, the learning rates were adjusted to compensate. For example, for the weights from feedforward to excitatory neurons and for recurrent excitatory weights:

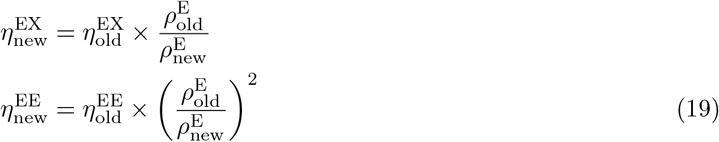

Additionally, simulations with different number of neurons were run. Multiples of the total number of neurons in the original model were chosen: *c*_n_ ∈ *{*1, 2, 3, 4*}*. For the input neurons, there would be *c*_n_ ON and OFF neurons per pixel, leading to *c*_n_*N* ^X^ input neurons; for the output excitatory and inhibitory neurons, there would be *c*_n_*N* ^E^ and *c*_n_*N* ^I^ neurons respectively. All-to-all plastic on feedforward and recurrent weights remains.

The simulation parameters used when varying total neuron numbers remain the same (Table 3) except for the number of neurons, L1 norms and learning rate. A condition that arises from balance theory is that synaptic weights scale as 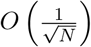 (Gu et al., 2018; Tian et al., 2020; van Vreeswijk and Sompolinsky, 1998), which has also been observed *in vitro* (Barral and D Reyes, 2016). Therefore, the L1 norms scale as 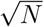. Additionally, the learning rates are adjusted as in Equation 18.

## 4 Results

### 4.1 Sparseness in the Network’s Activity

It is known that neurons in the sensory cortex have a sparse response to stimuli (Willmore and Tolhurst, 2001) with experimental results providing quantified assessment using the modified Treves-Rolls sparseness metric (Tolhurst et al., 2009).

To investigate this property the network was trained and the weights converged to stable values. The membrane potential plot of a representative neuron is shown in Figure 2A and the spiking activity of all neurons in the network in response to a single image is illustrated in Figure 2B. This spike data across images is converted into spike rates per second for each neuron in response to each image and is shown in Figure 2C for the ON, OFF, E, and I populations. For clarity of representation the response of only 30 neurons in each population are shown in response to only 30 images in the batch of 100. Each neuron in each population appears to be selective, i.e., responding strongly only to certain images (temporal sparseness).

**Figure 2:**
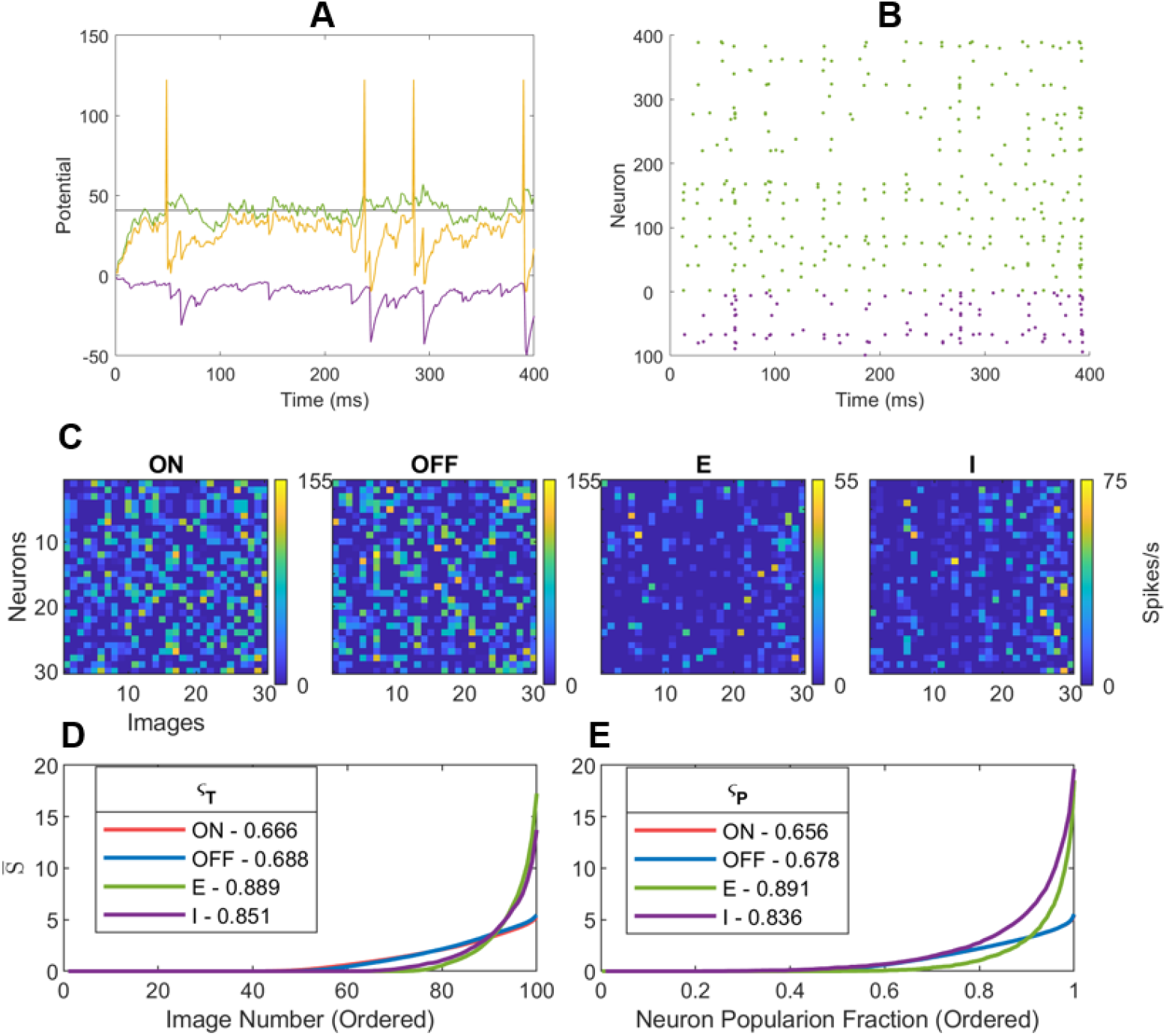
Sparse Network Response After Training: (A) The membrane voltage response of an excitatory neuron (E-388) in response to a single image presented for 400 ms after learning (green is from excitatory neurons, purple is from inhibitory neurons, yellow is net voltage, and black is the spiking threshold). (B) (A) The raster plot of the spiking response of all trained neurons (green is for 400 excitatory neurons and purple is for 100 inhibitory neurons) in response to a single image. The population sparseness is 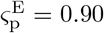 and 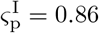 for the excitatory and inhibitory population respectively. (C) The spike rate of a subset of 30 neurons in each neural population (ON, OFF, E, and I) in response to a subset of 30 images. (D) Temporal Sparseness in response to 100 images. This is calculated by first sorting each neuron’s response (the rows of the arrays in (C)) from lowest to highest response spike rate before then averaging across rows to give mean spikes rate 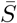 Note that the red and blue traces are closely aligned. (E) Population Sparseness. This is calculated by first sorting each image’s response (the columns of the arrays in (C)) from lowest to highest response spike rate before then averaging across columns. Neural parameters as described in Table 3

To quantify the temporal sparseness of the responses shown in Figures 2C each of the rows were ordered from lowest to highest response across the 100 images used, these ordered rows were then averaged together to give the average spike rate 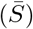 the curves shown in Figure 2D. This was done for each population ON, OFF, E, and I. The approximately exponential distribution of spike rates indicates that neurons were selective in their response, which is a feature indicative of spareness. The value of temporal sparseness for each population, 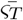 was calculated using Equation 16. The results are comparable to those of neurons found in the visual cortex (Tolhurst et al., 2009). The values of 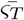 show that the E and I populations are more sparse than the ON and OFF populations and that the E population was more sparse than the I population.

As shown in Appendix B temporal sparseness in a LIF model is affected by the spike rate with a lower spike-rate leading to a higher temporal sparseness. This may explain the lower temporal sparseness of the inhibitory population.

Similar analysis for population sparseness is shown in Figure 2E with each of the columns in the arrays shown in Figures 2C ordered from lowest to highest response across all the neurons in each population. These ordered columns were then averaged together to give the curves shown in Figure 2E. The population sparseness metric, 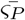 was calculated using Equation 17. The results are similar to those for temporal sparseness. To the extent that there is population sparsity this is related to decorrelation of temporally sparse neurons. If the response of a neuron is temporally sparse and it is decorrelated with other neurons in the population then this leads to population sparsity. This decorrelation is examined below.

### 4.2 Sparseness in the Network’s Receptive Fields

The spike triggered average (STA) for each neuron in the E and I populations was calculated, as described in the Methods subsection 3.9.1, and representative receptive fields are shown in Figure 3. A diverse range of receptive field shapes are learnt, such as localized unoriented blob-like filters and oriented Gabor-like filters. It can be seen that the network has learned Gabor-like receptive fields, which also arise from other sparse coding models (King et al., 2013; Lian et al., 2019; Olshausen and Field, 1997; Zylberberg et al., 2011). Furthermore, receptive fields of different sizes, positions and orientations can be seen for both excitatory and inhibitory neurons.

**Figure 3:**
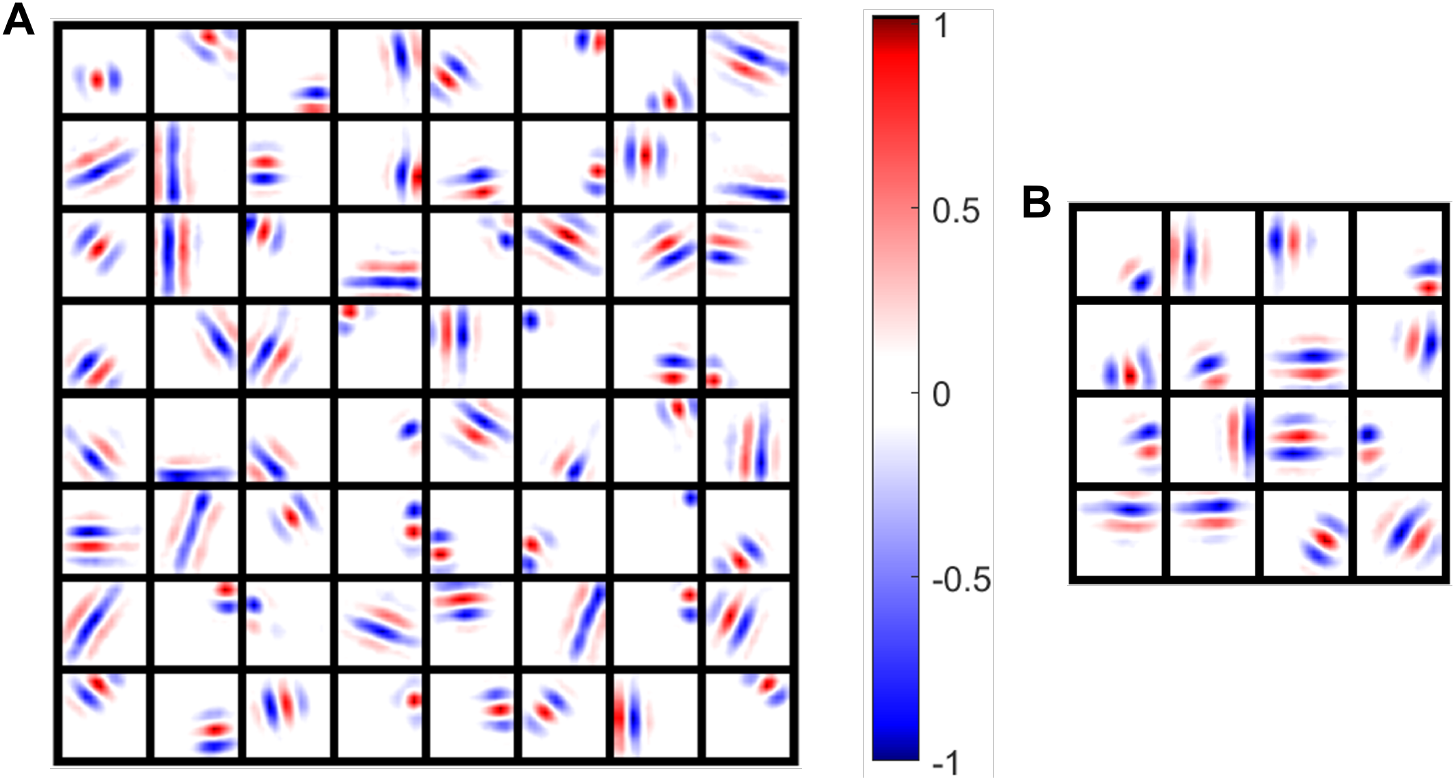
Receptive Fields of Excitatory and Inhibitory Neurons Characterized via Simulated STA: (A) Excitatory receptive fields of 64 randomly chosen neurons. (B) Inhibitory receptive fields of 25 randomly chosen neurons. Each box is a receptive field of a neuron where red represents ON and blue represents OFF which have values normalized. Neural parameters as described in Table 3.

These receptive fields were quantified by fitting each one to a parameterized Gabor function as described in the Methods subsection 3.9.1. The 369 out of 400 excitatory neurons that have a fit error less than 10% have been plotted with experimental data from cat and macaque monkey in Figure 4. The 31 cells that had RFs not well fit by Gabor filters tended to be in a transition state from one RF to another RF or have blob-like RFs. The resulting scatter plot has axes *n*_*x*_ and *n*_*y*_ which represent the length and width of the receptive field with respect to its spatial frequency. It can be seen that the Gabor functions are situated in the same region of parameter space as the Gabor functions measured from mammals *in vivo*. However, there are some regions in the *n*_*x*_, *n*_*y*_ space that is occupied by experimental results but not by the model. A parameter that affects the *n*_*x*_-*n*_*y*_ distribution is the target firing rate *ρ*^E^. Higher target firing rates lead to RFs that were more correlated and blob-like.

**Figure 4:**
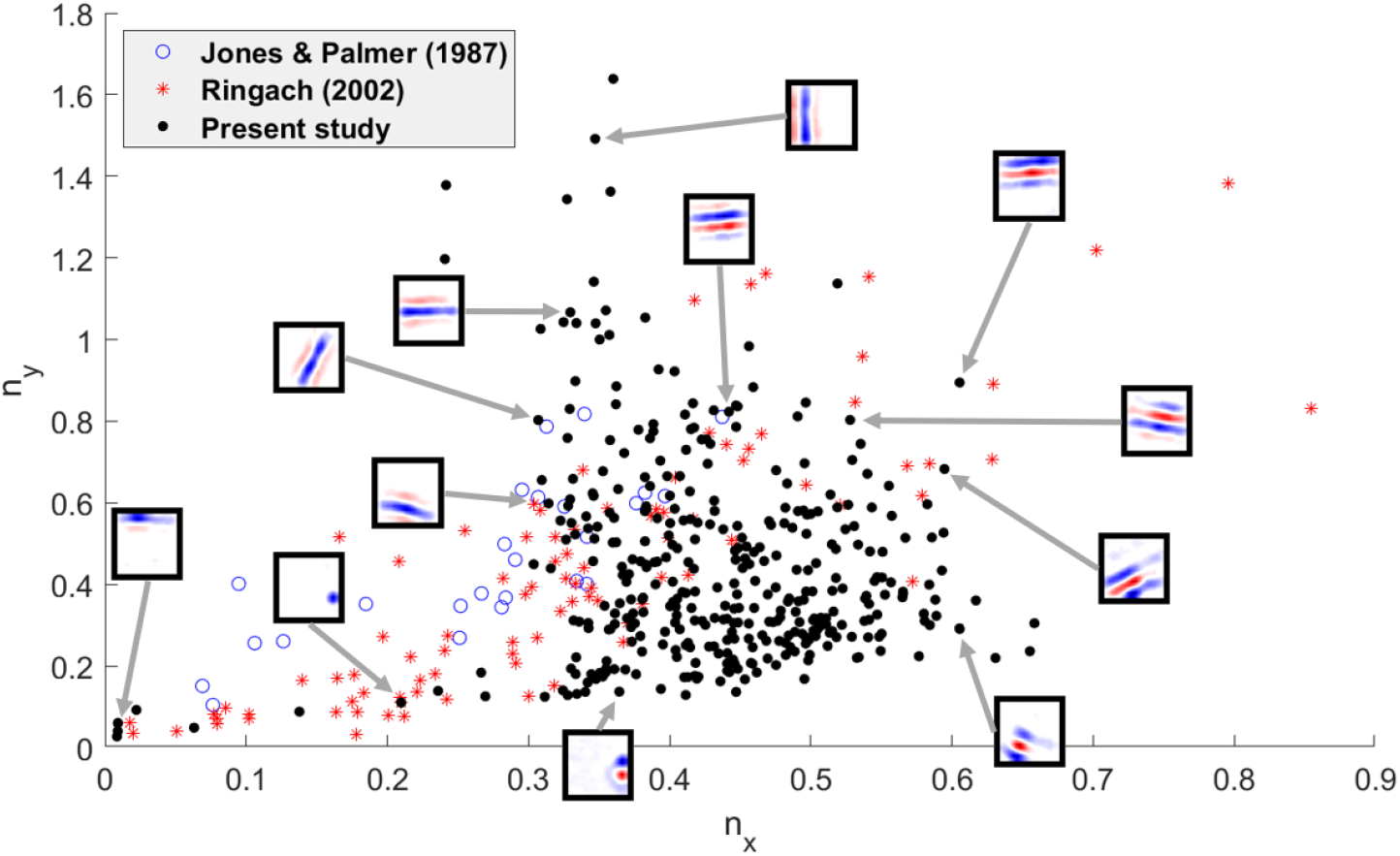
Comparison Between Receptive Field Shapes *n*_*x*_ vs *n*_*y*_: Distribution of receptive fields of model excitatory neurons compared with experimentally recorded receptive fields for cat (Jones and Palmer, 1987) and macaque monkey (Ringach, 2002). Open blue circles: cat data. Red stars: monkey data. Black dots: model data of 369 of 400 excitatory neurons that fit Gabor filters with *<*10% fit error. Example RFs are shown for model data.

### 4.3 Decorrelation in the Network’s Activity

To examine de-correlation of firing rates the spike rate data across neurons and images shown in Figure 2 was used. The correlation coefficient between the firing rates of each pair of LGN, E, and I neurons was calculated (Figure 5). The LGN and E populations are have small mean correlation coefficients and the distributions are also assymetric. In the LGN population this may be due to a bias in the statistics of the images used; a known feature of natural images (Ratliff et al., 2010). There appears to be an inhereted effect in the E population. The mean correlation coefficient between pairs of inhibitory neurons is higher. This increase in correlated firing for the I population is possibly due to the different feed-forward learning rules. The E population triplet rule discovers 2nd order correlations in the inputs (Brito and Gerstner, 2024), while the symmetrical rule used for the inhibitory population discovers 1st order correlations.

**Figure 5:**
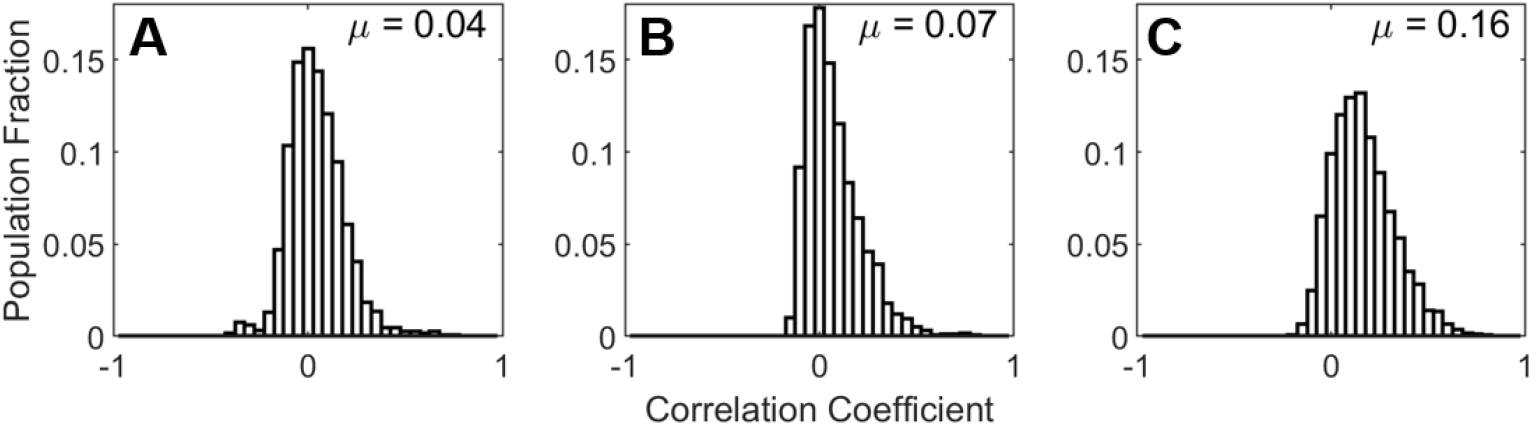
Decorrelation of Firing Rates using data from 100 images: (A) Pairwise correlation coefficients among the LGN population. (B) Pairwise correlation coefficients among the E population. (C) Pairwise correlation coefficients for the I population.

### 4.4 Decorrelation in the Network’s Receptive Fields

The receptive field of an excitatory neurons is shown in Figure 6A. The correlation coefficient of this neuron’s receptive field and each of the 100 inhibitory neuron’s receptive fields was calculated as described the the Methods, giving a distribution of values shown in the histogram in Figure 6B. The majority of the receptive fields cluster towards zero correlation while a minority have higher correlation or anti-correlation values. This is evidence of decorrelation between this particular neuron, and the population of inhibitory neurons.

**Figure 6:**
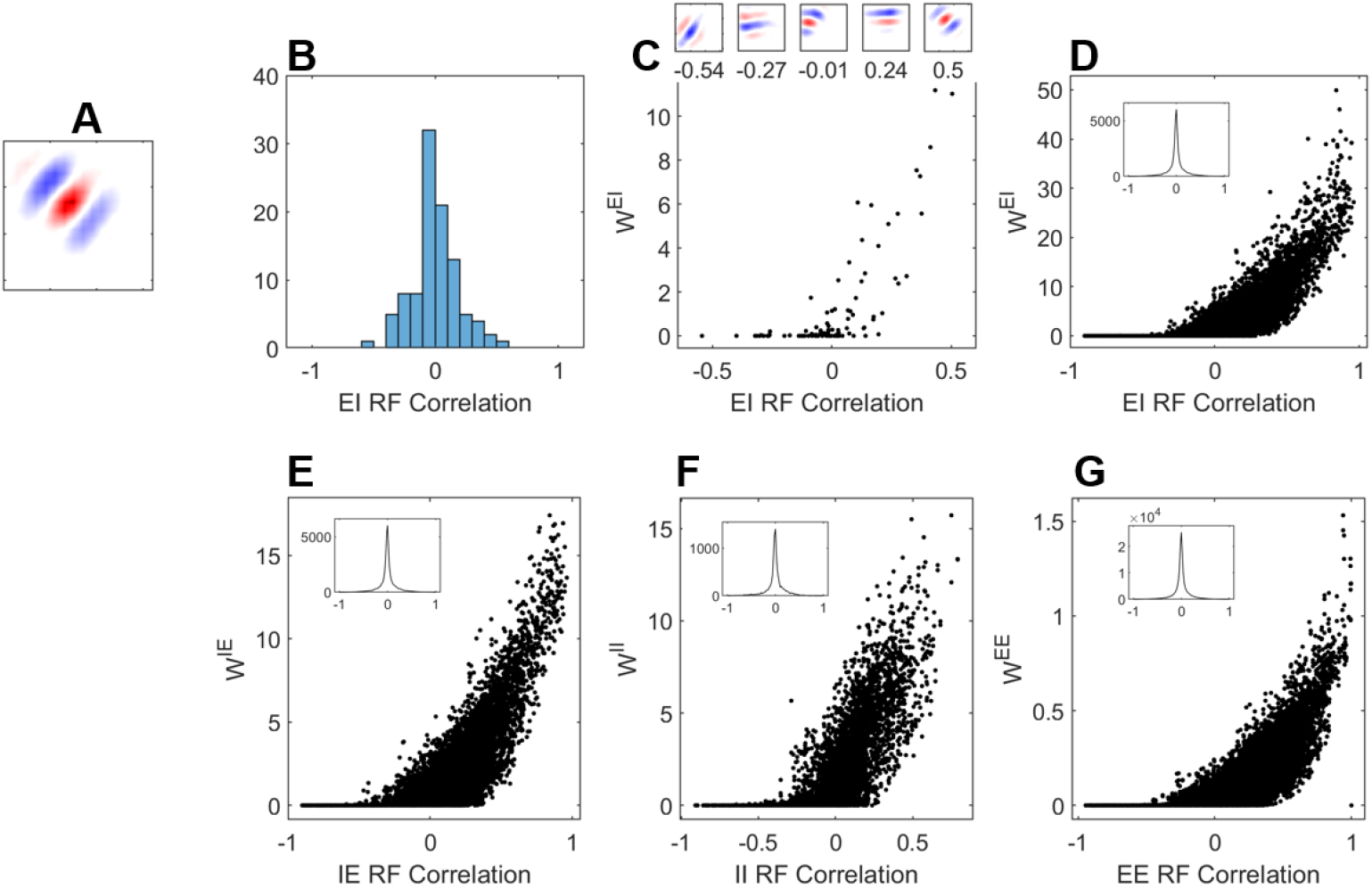
Decorrelation of Receptive Fields: (A) An excitatory neuron’s receptive field as measured via STA (B) The range of correlations between the this excitatory neuron’s receptive field and the receptive fields of the 100 inhibitory neurons in the network. (C) The inhibitory input weight from each of the inhibitory neurons as a function of their receptive field correlation. Example inhibitory neuron receptive fields at different correlation values are shown in inset. (D-G) The input weight to as a function of the pairwise receptive field correlations for every EI neuron pair, IE neuron pair, II neuron pair and EE neuron pair. The insets show the histogram of the RF correlations using bins of width 0.02. Neural parameters as described in Table 3.

The pairwise correlation coefficient was plotted for all 40,000 pairs of neurons (Figure 6C, inset). There is decorrelation between these two populations, with coefficients clustered at 0 and no apparent positive bias. The same analysis reveals similar uncorrelated receptive fields among all populations (Figure 6E-G, insets).

### 4.5 The Mechanism of Decorrelation in the Network’s Receptive Fields

To assess the role of inhibitory interneurons in decorrelating the responses of primary (excitatory) neurons the inhibitory input weight from all inhibitory neurons was plotted as a function of these correlation coefficients (Figure 6C). The same analysis was completed for all 40,000 pairs of inhibitory input to excitatory neurons in Figure 6D and for the reciprocal case of excitatory input to inhibitory neurons in Figure 6E. It can be seen that excitatory neurons receive strong input from inhibitory neurons with strongly correlated receptive fields (Figure 6D). Combining this with the fact that inhibitory neurons receive input from excitatory neurons with correlated receptive fields (Figure 6E) provides an indication of the underlying mechanism of decorrelation; excitatory neurons provide inhibitory input to other similar excitatory neurons via the inhibitory population, this is similar to the results in King et al., 2013. The resulting activity in the population of excitatory neurons will tend to avoid redundant representation of the visual input.

The data shown in Figure 6F shows that the inhibitory population has a similar decorrelation effect on other inhibitory neurons.

The data for excitatory-excitatory connections (Figure 6G) shows that there is a tendency for these weights to cause correlated activity in the network. In this network, the weights of excitatory to excitatory connections are low compared to other lateral weights, and so the effect is negligible. However, it is possible that this apparently detrimental mechanism has a beneficial role in temporal tasks where direction selective cells provide additional information, something not investigated here.

### 4.6 Balance in the Network’s Activity

Biological networks in the brain are known to exhibit balance in the contributions of excitation and inhibition to each neuron. The neuron in Figure 2A appears to be in a fluctuating regime (Destexhe et al., 2001; Meffin et al., 2004), where the membrane potential tends to sit just under the spiking threshold, and fluctuations cause irregular spiking activity. This appears to be due to a balance of excitation and inhibition (the green and yellow traces).

To examine this property in the trained network an excitatory neuron was randomly selected, whose receptive field as characterised by STA is shown in Figure 7A.

**Figure 7:**
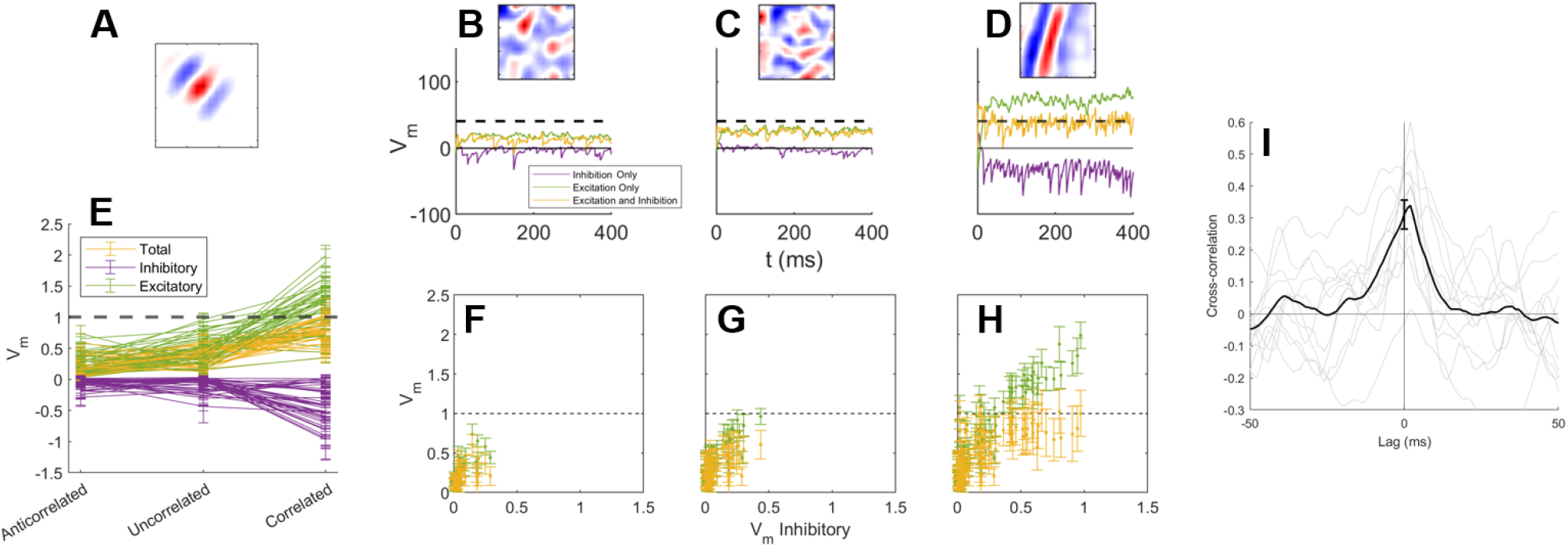
Loose Balance: (A) An excitatory neuron’s receptive field as measured via STA. (B) The neuron’s membrane voltage during the network’s response to the anti-correlated image stimulus shown in the inset. The image was selected as the most anti-correlated from 1000 images. The spike-threshold and voltage minimum was removed to allow observation of the full inhibitory response. The response without inhibition and without excitation is shown in green and purple. (C) Response to the most uncorrelated image. (D) Response to the most correlated image. (E) Using the same approach as in (B-D), the average membrane voltage for 40 randomly chosen neurons in response to the most anti-correlated, un-correlated, and correlated images (chosen from 100 random images). Error bars show the standard deviation in the voltage. (F-H) The excitatory and total membrane membrane voltages plotted as a function of the amplitude of the inhibitory only voltage for using the data in (E). (I) The cross correlation between the excitatory-only and inhibitory-only responses of the neuron in Figure 7 when input is provided by ten correlated images patches similar to that shown in Figure 7D (inset). The black trace shows the mean of the ten traces and the error bar gives the confidence interval of the mean at a lag of 0 ms. Neural parameters as described in Table 3.

To show that it receives balanced input it is necessary to compare the total influence of excitation and inhibition across images that the neuron responds strongly to and weakly to. To select input stimuli the whitened image patch patch correlation was calculated with the selected neuron’s receptive field. The most anti-correlated (low-input), uncorrelated, and correlated (high-input) image patches were chosen from random set of 1000 images. These three images provide the neuron with its lowest to highest input drive and are shown in Figure 7B-D (insets). To quantify the total excitatory and inhibitory input the neuron’s spike-threshold and minimum membrane voltage limit were both removed to allow the membrane voltage to evolve without interference.

The membrane voltage during the network’s response to these images is shown in Figure 7B-D (yellow trace). To allow comparison of excitation and inhibtion either the inhibitory or excitatory inputs were removed so that the membrane voltage was due only to either the excitatory or inhibitory input. With identical spiking input, the membrane voltage with only excitatory input and with only inhibitory input is also shown (Figure 7B-D, green and purple traces).

It can be seen that the increase in excitation during the network’s response to the correlated image patch was accompanied by an increase in inhibition, as shown in Figures 7E & G. The outcome was that the total membrane voltage, with both excitation and inhibition, had a mean and standard deviation that rarely transited above the spike-threshold, despite the increase in input, and was not excitation dominated.

This behavior was ubiquitous across the network as shown by a similar analysis of the mean voltages across 40 neurons (Figure 7E). To examine the presence of balance in more detail, for these 40 neurons the mean excitatory input was plotted as a function of the mean inhibitory input for low to high input levels (Figure 7F-H, green) showing a correlation. Most convincingly, plotting the total mean membrane voltage (yellow) as a function of the mean inhibitory input shows that the total voltage is brought down below the spike-threshold ensuring that the neurons is not excitation dominated, even at high input levels. This is most apparent in Figure 6 H).

Tight balance is associated with transient modulations in the excitatory and inhibitory inputs, as illustrated in Figure 1E. To examine this property, the cross-correlation between the excitatory and inhibitory response to the ten high input image patches was calculated as described in the Methods (Figure 7 I). The period of response from 0 to 100 ms was excluded to avoid measurement of correlations in the initial transient change in membrane voltage.

Figure 7 I shows that there was a very high degree of correlation of the excitatory and inhibitory inputs to the neuron. The correlation decays with a time-window of approximately 10 ms, a similar time-scale to that of the membrane time constant, *τ*_m_. This fast temporal correlation on the time scale of *∼*10 ms is an example of tight balance. It can also be seen that the inhibitory input lags the excitatory input by approximately 2-3 ms. This delay is likely to be a combination of the extra synapse and the integration of inputs by the inhibitory neurons.

### 4.7 Robustness of the Network

#### 4.7.1 Scaled Target Firing Rates

When the target firing rate is changed, metrics such as excitatory receptive field correlation, temporal and population sparseness are affected, as illustrated in Figure 8A-C. Excitatory receptive fields become more correlated with each other when their target firing rate is increased. With a higher firing rate, more neurons tend to be active with a given stimulus. As a result of overlapping activity, their receptive fields would have some overlap as well, as shown in Figure 8A. An increased target firing rate leads to a decrease in temporal sparseness, as shown in Figure 8B. The reason for this is explored in Supplementary Material Section 6. As temporal sparseness decreases it is expected that this leads to population sparseness decrease, and this is shown in Figure 8C.

**Figure 8:**
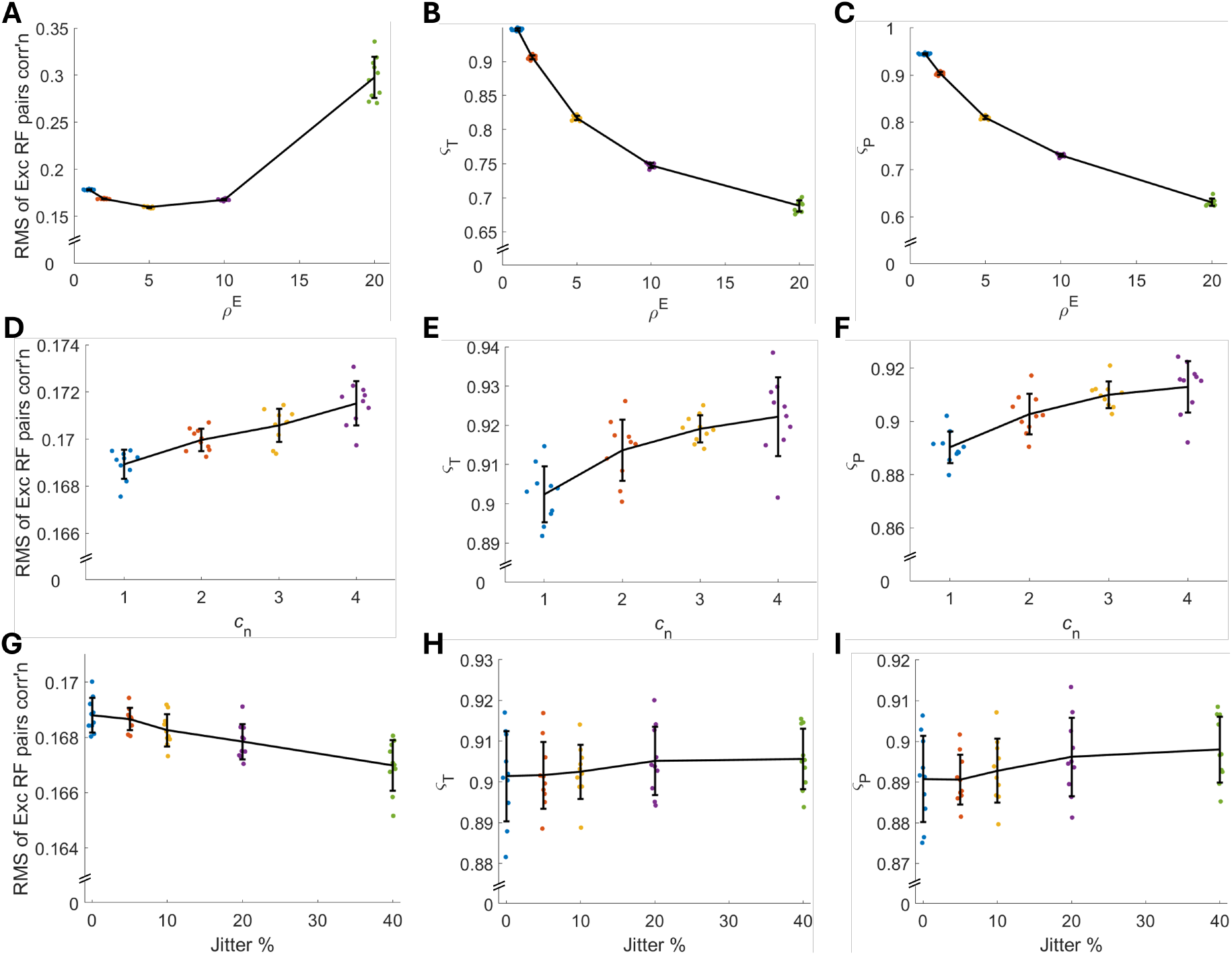
Robustness of the network: (A-C) Firing rates of excitatory and inhibitory neurons are varied. (D-F) Total number of neurons in the network are varied, where *c*_n_ is the multiple of the default total number of neurons. (G-I) Mean weights are jittered. (A,D,G) RMS of the Pearson correlation of the receptive fields between all excitatory cell pairs. (B,E,H) Temporal sparseness, *ς*_T_, is measured. (C,F,I) Population sparseness, *ς*_P_, is measured. Neural parameters are varied as described in Methods 3.9.5 and 3.9.6. Other neural parameters as described in Table 3. Black lines connect the means, and errorbars display the standard deviation of individual simulations represented as coloured dots that have a random jitter in the x-axis for visibility.

#### 4.7.2 Scaled Network Size

When the total number of neurons increases, the metrics of excitatory receptive field correlation, temporal and population sparseness all remain relatively constant, as shown in Figure 8D-F. This demonstrates that the network can scale up to larger neuron numbers, as seen physiologically. It is important to note that when scaling up the network, the mean weight value was adjusted to scale as 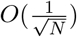, as is observed physiologically (Barral and D Reyes, 2016).

#### 4.7.3 Jittered Paramaters

When the L1 weight norms are varied up to 40%, metrics such as excitatory receptive field correlation, temporal and population sparseness remain relatively constant, as shown in Figure 8G-I. Simulations with different mean weight values would result in different amounts of excitatory and inhibitory inputs which could potentially cause an excitatory-inhibitory imbalance. However, all simulations conducted have resulted in stable learning and activity. One reason is the adaptive spiking threshold, which is adjusted to ensure that neurons fire within stable levels. As well as remaining stable, certain metrics remain at similar values despite changes in the weight norms. This includes the pairwise excitatory receptive field correlation, temporal and population sparseness.

## 5 Discussion

In this study a spiking network model of V1 was developed using a bottom-up approach based upon biological principles, including separate ON and OFF inputs, spiking neurons, separate excitatory and inhibitory populations (Dale’s Law), spike-timing-dependent plasticity, and firing rate homeostasis. It was found that after training the model exhibited several properties observed in biological systems: a sparse neural responses, decorrelation of activity and receptive fields, and balanced excitation and inhibition. Additionally, the network was robust to changes in network size as well as to random jitters to the synaptic weights, but not to high target firing rates of the output neurons.

### 5.1 Emergence of Sparseness in the Network

The sparse *temporal* spiking activity observed in Figure 2C and quantified in Figure 2D appeared to be similar to available experimental data (Tolhurst et al., 2009; Yoshida and Ohki, 2020). These values were however shown to be vulnerable to increases in target spike rate (Figure 8B). Using analysis of the LIF model (See:Supplementary Material S3) this was discovered to be a feature of the LIF model; without other changes, temporal sparseness decreases with increasing spike rate. Other features of the LIF neuron also influence sparseness such as the synaptic and membrane time constants. It is unknown whether this is also a feature of biological cortical neural responses, but is a possible avenue for future experimental work.

This temporal sparseness is known via mathematical sparse-coding models to be associated with learning of Gabor filters in the visual cortex. This is something we also found in our model (Figures 3 and 4).

Experimental measures of population sparseness are difficult to obtain because they simultaneous measurement from multiple cortical neurons. However the sparse *population* spiking activity observed in Figure 2C and quantified in Figure 2E appears to qualitatively match the available experimental data (Yoshida and Ohki, 2020).

These collection of results indicate that the network has sparse properties without sparseness being achieved via an objective function or otherwise explicitly imposed. Instead, the LIF model with adaptive excitability established a sparse firing response to stimuli which provided correlations for the triplet-STDP rule to discover.

### 5.2 Emergence of Decorrelation in the Network

Figure 5 shows that the mean spike rate correlation coefficient was positive for the excitatory (Figure 5B) and inhibitory populations (Figure 5C). It is unknown whether this non-zero mean population decorrelation is a feature of biological cortical neurons and this perhaps is a question for experimental work. This appears to be an effect that is inherited and amplified from the LGN inputs which also showed a positive mean correlation coefficient (Figure 5A). LGN neurons are known to show a bias towards OFF features in natural scenes (Ratliff et al., 2010).

The decorrelation of receptive fields within the E (Figure 6G inset) and I (Figure 6F inset) populations did not reflect this positive bias in spike response correlations. Instead the receptive fields were well de-correlated.

These results showing decorrelation of activity and receptive fields were again, not something that was imposed explicitly using an objective function. Instead, symmetric STDP rules in inhibitory neurons discovered correlated firing in excitatory neurons (Figure 6E. Effectively, they are gathering evidence that those excitatory neurons have similar receptive fields. Simultaneously, symmetric STDP rules in excitatory neurons discover correlated firing in inhibitory neurons (Figure 6E gathering evidence of their similar receptive fields. Combined, this leads to indirect inhibition between excitatory neurons (King et al., 2013).

### 5.3 Emergence of Balance in the Network

Correlations in the magnitude of total excitatory and inhibitory input to excitatory neurons were observed in the trained network (Figure 7D, 7E, and 7H). These ensured that even with strong input, the neurons did not become excitation dominated. There was also an associated fine temporal correlation between these excitatory and inhibitory inputs (Figure 7I).

The total excitatory and inhibitory input to each neuron is decided by balance network theory using the inequalities in Equation 11 enforced via the homeostasis described by Equation 9. However, this is insufficient to ensure the resulting tuned and fine balance in Figure 7. Instead this balance has emerged as a consequence of a combination of the homeostasis and the synaptic learning rules that ensure that excitatory neurons receive input from inhibitory neurons with similar receptive fields.

### 5.4 Relationship between Sparseness, Decorrelation, and Balance

Population sparseness in activity is partly the result of decorrelation due to lateral synaptic plasticity. The population sparseness in activity results in changes in the correlations discovered by the excitatory neurons among their feed-forward input weights from LGN neurons. Once these changes in receptive field have taken place, population sparseness in activity is also due simply to the decorrelation in receptive fields among the E neuron population. In this way population sparseness and decorrelation operate on the fast time scales of the network’s immediate response (*∼*10 ms) to inputs but also the slow time scales of synaptic plasticity (*∼*1 h).

This population sparseness in excitatory neuron receptive fields ensures that the network maximizes the information it carries about its inputs. This is something that an ‘infomax’ function would normally explicitly require to ensure statistical independence (Bell and Sejnowski, 1995; Linsker, 1988). In this BGNN emerges from learning rules governing the connections from inhibitory neurons.

AS highlighted above, the tight balance shown in Figure 7I is closely related to decorrelation. The correlated inhibitory inputs prevents the neuron from firing at times that are are predictable by other neurons in the network. This suppression of predictable responses is reminiscent of the principles of efficient coding (Barlow, 1961), and predictive coding (Rao and Ballard, 1999).

### 5.5 Comparison of Receptive Fields with Experimental Studies

The network model in this study has separate excitatory and inhibitory populations with all-to-all feed-forward and recurrent connectivity that are learnt via STDP. When whitened images were provided to the LGN input neurons, the output neurons learned visual receptive fields that resembled the shape and diversity of receptive fields measured experimentally, as illustrated in Figure 4. In particular, localized un-oriented blob-like receptive fields as well as localized oriented Gabor-like receptive fields of varying sizes and spatial frequencies were observed to closely resemble those observed in physiological studies (Ringach, 2002). Inhibitory neurons show a range of properties in experimental studies (Markram et al., 2004), including orientation tuning, and they have receptive fields that resemble those of excitatory cells (Hirsch and Martinez, 2006; Hirsch et al., 2003). The inhibitory neurons in the network learned receptive fields that resemble those found in experimental studies.

### 5.6 Spike Timing Dependent Plasticity

The synaptic plasticity rules used in the study presented here are those that have been observed experimentally. In particular, the type of STDP implemented depends upon the identity of the presynaptic and postsynaptic neuron. Synapses where the presynaptic and postsynaptic neuron are both excitatory neurons learn via the STDP triplet rule (Pfister and Gerstner, 2006). This rule fits experimental data well and can be mapped to the BCM rule in rate-based neural models, which has been used to model visual phenomena (Cooper and Bear, 2012). A recent spiking network used the triplet rule to also produce visual receptive fields and showed that using the triplet rule allowed their neurons to ignore low-order correlations and find features hidden in higher-order statistics (Brito and Gerstner, 2024). Their model, however, implements an output population only with one output population with recurrent inhibition, instead of separate excitatory and inhibitory subpopulations. In our model, inhibitory synapses learn through symmetrical STDP. Experimental studies have observed that the order of spike times (pre-before-post or vice versa) does not affect inhibitory plasticity (D’amour and Froemke, 2015). Moreover, symmetrical STDP can be mapped to rate-based Hebbian learning (Vogels et al., 2011, Supplementary Material). The results here indicate that symmetrical STDP, because of its associative Hebbian-like learning, is sufficient to find correlations between the output excitatory and inhibitory neurons through their recurrent connections, similar to the correlation-measuring rule used in King et al. (2013). With the inhibitory population, this rule acts to de-correlate responses and receptive fields.

### 5.7 Homeostasis

The STDP rules that lead to the network structure and selectivity in this study are separate from the homeostatic processes, which maintain stability and balance. Biological homeostasis typically operates on a slower timescale of hours or days (Turrigiano, 2008; Watt and Desai, 2010). If only STDP and Hebbian plasticity were present, this would lead to pathological runaway dynamics (Abbott and Nelson, 2000). Therefore, there is a requirement for compensatory and stabilizing synaptic processes in computational models (Zenke and Gerstner, 2017), typically implemented by homeostatic synaptic processes (Turrigiano, 2008). In the network, there are two homeostatic processes: weight normalization (Turrigiano, 2012), and an adaptive spiking threshold (Zylberberg et al., 2011), which can be likened to activity-dependent synaptic scaling and intrinsic plasticity respectively (Debanne et al., 2019; Turrigiano, 2012; Turrigiano et al., 1998. Both of these homeostatic mechanisms directly ensure that the weights and the firing rates, respectively, remain stable. In the study, these processes were found to be sufficient for network stability and function: when the mean weight values for the different connection groups were jittered randomly, or when the total number of neurons were scaled up, the network behaviour remained robust to perturbations of the network parameters, and the network was still able to learn the diverse range of V1 RFs.

### 5.8 Future Work

In this study, current-based synapses were used, where synaptic inputs were treated as currents directly injected into the neuron. However, using conductance-based synapses would be more biologically realistic. Conductance-based synapses would allow the effective time constant to change to maintain excitatory and inhibitory balance over a wide range of firing rates (Burkitt, 2006). Consequently, this may affect temporal and population sparseness.

The network included a lateral excitatory-excitatory pathway, which was given a low strength. The functional role of lateral excitatory connections in the brain remains unclear but is likely to be associated with the representation of temporal features of visual information that were not explored in this study. Increased weight of excitatory-excitatory connections can easily lead to pathological spiking. The use of spatiotemporal stimulus, such as natural video is therefore an important aspect of future work.

## 6 Acknowledgements

This work was supported by the Australian Government through the Australian Research Council’s Discovery Projects funding scheme [DP220101166].

## S1. Network with weights initialized to low values learnt biological receptive fields

The simulations presented in the main text had the weights initialized to the L1 norm upper bound. This section address the question of whether the network is able to learn appropriate receptive fields when the weights are initialized to low values. The weights are initialized as described in Methods 3.6, with one difference: the initial L1 norm of the weights was set to 1. After learning, the L1 norms of the weights reached their respective L1 norm upper bounds, and appropriate receptive fields are observed as seen in Figure S1A for excitatory neurons and Figure S1B for inhibitory neurons. These receptive fields are qualitatively similar to the receptive fields in the main text (Figure 3).

## S2. Baseline firing before and after learning

Before presenting natural images, the weights were kept fixed and the input neurons fired at a background rate of 1 Hz. The spiking thresholds were allowed to adapt and the target firing rates of the excitatory and inhibitory neurons were set to 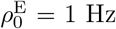 and 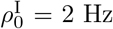 respectively. This represents the resting state of the network before learning, where all neurons fire with a spontaneous activity. Following this, the target firing rates become *ρ*^E^ = 2 Hz and *ρ*^I^ = 4 Hz respectively when presented with static visual stimuli.

The synaptic weights in the network of spiking neurons described here changed due to learning with static image stimuli as described in Subsection 3.4. Neural circuits need to maintain plasticity to accommodate changes in connectivity and synaptic strength during development and learning. Also important is maintaining the spontaneous firing of neurons before and after learning for the stability and function of neural circuits. However, it is a challenge to maintain both ongoing plasticity and stability at the same time (Schulz, 2006). Baseline activity is maintained throughout the learning process as a result of synaptic scaling implemented by L1 weight normalisation (as described in Subsubsection 3.4.3).

Prior to the learning process, when the network is presented with background firing rates, where all input neurons fire at the same spontaneous rates, the output neurons fire at some level proportional to this input rate. After the learning process using natural images, when the spiking thresholds were returned to the levels before learning through a homeostatic mechanism for the output firing rate (Turrigiano, 2012), the input-output response curve appears to be similar before and after learning, as shown by comparing Figure S2A with Figure S2B. However, the input-output response curve before learning is determined by the initial conditions and the method of weight initialization, while the input-out response curve after learning is determined by the learnt weights that underwent plasticity.

## S3. The effect of Target Rate on Sparseness

To show why temporal sparseness (*ς*_T_) decreases with increasing target spike rate (*ρ*^E^) it is possible to use Siegert’s formula for spike rate of a LIF neuron (Burkitt, 2006):

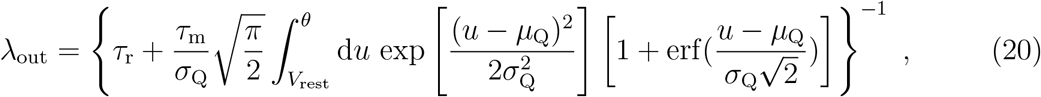

where *τ*_*r*_ is the absolute refractory period, *τ*_m_ is the membrane time constant, *V*_rest_ is the reset membrane voltage and *θ* is the spike threshold voltage. *µ*_Q_ and *ς*_Q_ are the mean and variance of the free membrane potential with *δ*-current input from an excitatory Poisson process and inhibitory Poisson process:

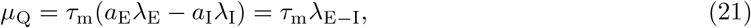

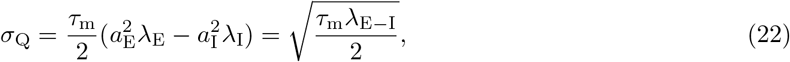

where *a*_E_ is the amplitude of the excitatory input synapse, and *λ*_E_ and *λ*_I_ are the excitatory and inhibitory Poisson input rates. The simplified formulations of equations 21 and 22 are derived by substituting *a*_E_ = *a*_I_ = 1 and *λ*_E*−*I_ = *λ*_E_ *− λ*_I_.

Using equation 20 with *a*_E_ = 1, *τ*_m_ = 10 ms, *τ*_r_ = 1 ms, and a reset voltage *V*_rest_ = 0 the output rate was calculated for 100 input rates (*λ*_E*−*I_) logarithmically spaced between 10 Hz and 1000 Hz. This is a simulation of the statistics across images of the high-weight input synapses. The value of *θ* was adjusted until the average output rate 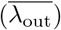 across all these inputs was 1, 10, or 100 Hz. The results are shown in Figure S3A.

It is clear that a lower target spike rate, associated with a higher threshold, gives lower spike rates in response to the range of input spike rates. However, by plotting these rates as a multiple of the mean it can be seen that the lower target rate appears to give a significantly more sparse distribution of output rates (Figure S3B). This is confirmed using the Treves-Rolls sparseness metric (Equation 16). The LIF neuron with a threhsold adjusted to achieve a 1 Hz mean output rate gave a value of *ς*_*T*_ = 0.94 while the 10 Hz

**Figure S1:**
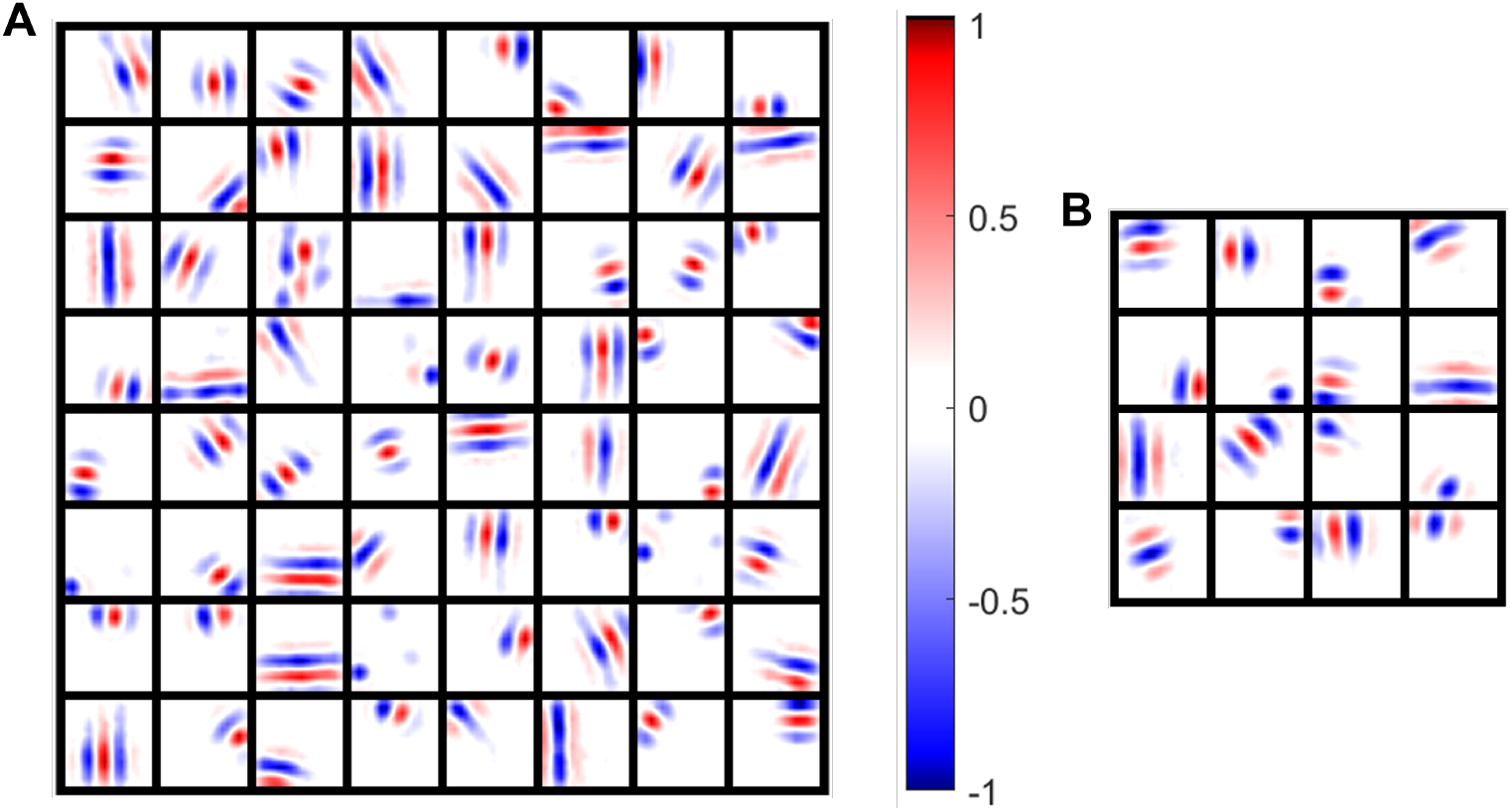
Receptive Fields of Excitatory and Inhibitory Neurons when Small Weights are Initialised: (A) Excitatory receptive fields of 64 randomly chosen neurons. (B) Inhibitory receptive fields of 25 randomly chosen neurons. Each box is a receptive field of a neuron where red represents ON and blue represents OFF which have values normalized. Neural parameters as described in Table 3.

**Figure S2:**
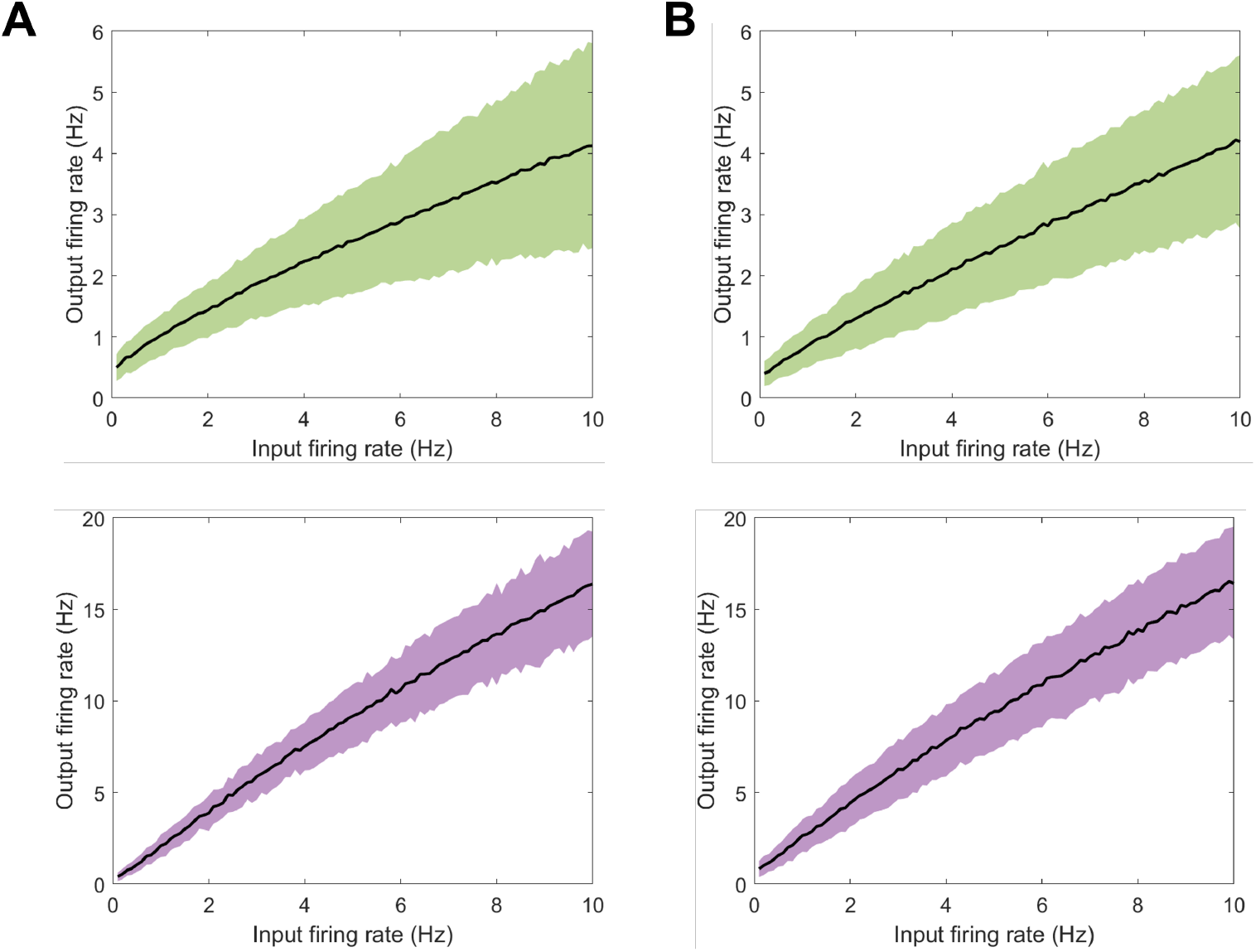
Network Response to Spontaneous Input: Response of output neurons is plotted when input neurons have a constant firing rate (A) with initialised weights before learning and (B) after learning with natural images with spiking thresholds set back to the same level as before learning. Mean (black line) and standard deviation (blue for excitatory and red for inhibitory neurons) are plotted. Neural parameters as described in Table 3. mean output rate resulted in *ς*_*T*_ = 0.59. Plotting these values results in a curve (Figure S3C) that can be compared to the trend observed in the full spiking model (8B).

**Figure S3:**
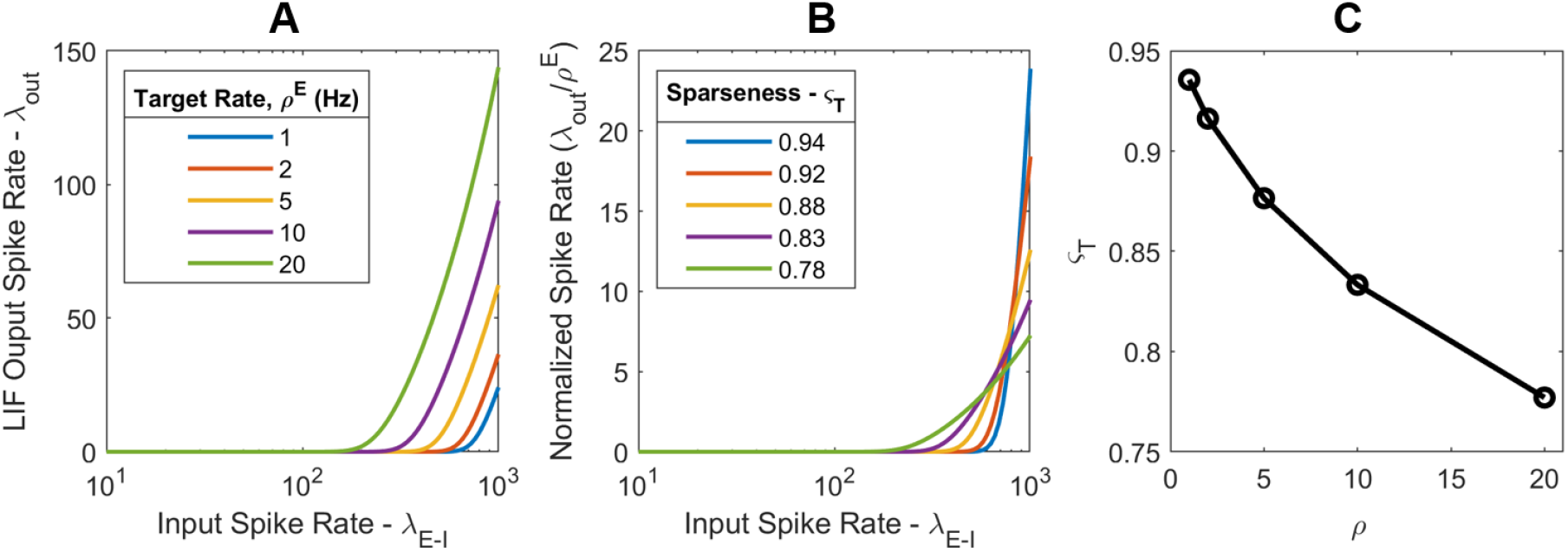
Spike Rate and Sparseness: (A) Output spike rate of a LIF neuron as a function of excitatory Poisson input rate. The spike-threshold has been adjusted to produce a mean spike rate across all inputs of 1, 10, or 100 Hz (B) The same data normalized to the target spike rates with the sparseness metric now shown in the legend. (C) The resulting calculated sparseness values as a function of the target rate.

## Notes

https://github.com/marko-ruslim/bio-spiking-v1-learning

### Competing Interest Statement

The authors have declared no competing interest.

